# Alkoxylated fisetin derivatives – the new potential drugs against head and neck cancers

**DOI:** 10.64898/2026.02.03.703451

**Authors:** Chudy Patryk, Wala Paulina, Krzykawski Kamil, Kadela-Tomanek Monika, Dziedzic Arkadiusz, Kubina Robert

## Abstract

Head and neck squamous cell carcinoma (HNSCC) is a therapeutically challenging cancer what underscoring the need for new chemical agents that selectively induce programmed cell death. Fisetin, a naturally occurring flavonoid, exhibits promising anticancer activity but displays limited proapoptotic efficacy and selectivity. Here, we examined whether alkoxylated modification of fisetin enhances its ability to induce apoptosis in HNSCC cells.

Fisetin derivatives bearing four-carbon substituents were synthesized and evaluated in multiple HNSCC cell lines. Two derivatives, MKT218 and MKT257, markedly reduced HNSCC cell viability at low micromolar concentrations with low toxicity towards normal human fibroblasts. Notably, the observed cytotoxicity was not associated with activation of a canonical DNA damage response, as neither γH2AX accumulation nor p53 activation was detected. Furthermore, PARP1 cleavage and live-cell imaging combined with annexin V/EthD-III staining revealed a significantly higher proportion of apoptotic cells. The effect was stronger following treatment with MKT218 and MKT257 compared with fisetin. Time-lapse microscopy further demonstrated that fisetin derivatives, particularly MKT218, promote mitosis-associated apoptosis, in contrast to the predominantly cytostatic effect of fisetin. Moreover, *in silico* docking suggested that MKT218 exerts its pro-apoptotic activity through a multi-target interaction profile involving key regulators of cell survival and apoptosis rather than a single dominant target.

To sum up, our findings suggest that alkoxylated fisetin derivatives may be constituted as new non-genotoxic inducers of apoptosis in HNSCC cells.

## 1. Introduction

Head and neck cancer (HNC) is a group of malignancies that are placed mainly in the lips, oral cavity, oropharynx, larynx and salivary glands, with 90% of tumours diagnosed as head and neck squamous cell carcinoma (HNSCC). HNC has been identified as the 7^th^ most common cancer worldwide, with almost 1 million new cases and 0.5 million deaths worldwide in 2022 [1, 2]. The main risk factors driving the neoplastic development of the HNC are tobacco use (smoking and chewing), alcohol consumption, low socio-economic status and human papilloma virus (HPV) infection, with the last affecting the therapeutic approach [3, 4]. The diagnosis is also more common among men and people at the age of 50-70 years [5]. Due to the divergence of anatomical sites affected by HNC, the symptoms are not unified and are specific to the affected tissue, making the diagnostic process difficult. Regarding the difference between HPV-positive and HPV-negative HNC, the statistical analyses show that HPV-positive cancers are more frequent in the oropharyngeal area, with rare diagnoses in the larynx or oral cavity [5, 6]. HPV-negative HNC has been associated with unfavourable prognosis, requiring extensive molecular diagnostics to tailor the treatment regime alongside the gold-standard use of cisplatin or 5-florouracil, while HPV-positive cancers are quite sensitive to radiation and anticancer drugs, which come with better prognosis [5, 7].

The general first line therapeutic strategy contain remains chemotherapy with the use of cisplatin and/or 5-florouracil, alongside surgical removal of the tumour and radiation. However, this approach presents several important challenges that limit its overall effectiveness. First, the immunisation of cancer cells can reduce the therapeutic response, making the treatment less effective over time. Second, chemotherapy is associated with numerous adverse side effects, including nephrotoxicity, gastrointestinal toxicity, and neurotoxicity. These complications place a significant burden on patients and substantially reduce their quality of life. Immunotherapy has also been found as an emerging therapeutical approach battling HNC, with positive outcomes post-treatment with pembrolizumab adjacent to chemotherapy. Similarly, targeted therapies have been growing in research extensively, specifically cyclin-dependent kinase inhibitors (CDKIs) and Poly(ADP-Ribose) Polymerase 1 (PARP-1) inhibitors. These medications have been found to work excellently in adjuvant therapies with chemotherapy, often resulting in synergistic effects, improving overall outcomes and decreasing systemic toxicity in patients [7].

Nonetheless, the risk of HNSCC resistance acquisition is inevitable and taken into account while designing the therapy course. Additionally, current therapies still pose risks of other syndromes induced by systemic toxicity. For this reason, research has been shifting onto natural compounds that have potential anti-cancer effects. Among these compounds are flavonoids, organic compounds naturally occurring in fruits and vegetables, known for their extensive positive effects in the organisms, ranging from anti-oxidative effects, neuroprotective, anti-aging, as well as anti-proliferative, anti-angiogenic and anti-mutagenic [8, 9]. Among these flavonoids, fisetin has been emerging as a potentially anti-cancer compound without cytotoxic effects on normal, healthy cells [9–12]. Fisetin has been found to promote cell cycle arrest in oral squamous carcinoma [13], bladder cancer [14], triple-negative breast cancer [15] and colon cancer [16] among many other cancer types [12]. Available data also suggest that fisetin significantly inhibits tumour growth breast cancer xenograft models, as well as increase the percentage of apoptotic cells when compared with control [17]. However, fisetin is hydrophobic, which results in poor systemic circulation and poor target towards tumour *in vivo* [17]. This limitation has inspired a search for advanced molecules of natural basis, resulting in synthesis of various flavonoid-derivative compounds that have substituted hydroxyl groups that exhibit stronger anti-cancer activity than the original compounds [18, 19]. In this paper, alcoxylic derivatives of fisetin have been synthesised and their *in vitro* activity has been evaluated to see if such modifications enhance the cytotoxic effect of the original compound on HNSCC cell lines.

## 2. Methods

### General synthesis of compounds

Fisetin (0.1 g; 0.350 mmol) and potassium carbonate (5 eq; 1.75 mmol; 0.241 g) were dissolved in acetone (4 mL). The 3-bromo-1-pronyne, 1-bromo-2-butyne or 1-bromobutane (5eq; 1.75 mmol) was added drop-wise and the mixture was stirred at reflux. The reaction progress was monitored using the thin layer chromatography (TLC) method. After reaction completed, the crude product was purified by column chromatography (SiO_2_, chloroform/ethanol, 40:1, v/v) to give pure compound:

∘ *2-(3,4-bis(prop-2-yn-1-yloxy)phenyl)-3,7-bis(prop-2-yn-1-yloxy)-4H-chromen-4-one* ***MKT218***: Yield 64%; mp. 159-160°C. The spectral data were confirmed by the literature [18].
∘ *2-(3,4-bis(but-2-yn-1-yloxy)phenyl)-3,7-bis(but-2-yn-1-yloxy)-4H-chromen-4-one* ***MKT257***: Yield 68%; mp. 173-174 °C; ^1^H NMR (600 MHz, aceton-d6) δ, ppm: 1.73 (t, J=2.4 Hz, 3H, CH_3_), 1.87 (t, J=2.4 Hz, 6H, 2xCH_3_), 1.88 (t, J=2.4 Hz, 3H, CH_3_), 4.85 (m, 2H, OCH_2_), 4.87 (m, 2H, OCH_2_), 4.93 (m, 2H, OCH_2_), 4.99 m, 2H, OCH_2_), 7.10 (dd, J=9.0 Hz, J=2.4 Hz, 1H), 7.25 (m, 2H), 7.84 (dd, J=8.4 Hz, J=2.4 Hz, 1H), 8.03 (d, J=1.8 Hz, 1H), 8.08 (d, J=9.0 Hz, 1H) (**Fig. S1a);** ^13^C NMR (150 MHz, aceton-d6) δ, ppm: 2.4, 2.5, 56.6, 56.7, 57.0, 59.2, 73.9, 74.0, 74.1, 74.2, 83.5, 83.6, 84.0, 84.1, 101.2, 113.4, 114.4, 115.1, 118.2, 122.4, 124.1, 126.6, 138.0, 147.2, 150.0, 155.3, 156.7, 162.2, 173.4 **(Fig. S1b);** IR (ν_max_ cm^-1^, ATR): 2963-2862, 2236, 1619, 1594, 1509, 1428, 1364, 1251; ESI-HRMS *m/z* [M+1]^+^ calcd for C_31_H_27_O_6_^+^ 495.1808, found 495.1809.
∘ *3,7-dibutoxy)-2-(3,4-dibutoxy)phenyl)-4H-chromen-4-one* ***MKT261***: Yield 62%; mp. 95-96 °C; ^1^H NMR (600 MHz, aceton-d6) δ, ppm: 0.93 (t, J=7.2 Hz, 3H, CH_3_), 1.01 (m, 9H, CH_3_), 1.46 (m, 2H, <u>CH_2_</u>CH_3_), 1.57 (m, 6H, <u>CH_2_</u>CH_3_), 1.72 (m, 2H, OCH_2_<u>CH_2_</u>), 1.83 (m, 6H, 3xOCH_2_<u>CH_2_</u>), 4.12 (m, 6H, 3xOCH_2_), 4.18 (t, J=6.0 Hz, 2H, OCH_2_), 7.05 (dd, J=9.0 Hz, J=2.4 Hz, 1H), 7.15 (m, 2H), 7.76 (dd, J=8.4 Hz, J=2.4 Hz, 1H), 7.82 (d, J=1.8 Hz, 1H), 8.05 (d, J=9.0 Hz, 1H) (**Fig. S2a)**; ^13^C NMR (150 MHz, aceton-d6) δ, ppm: 13.2, 13.2, 13.3, 18.9, 19.0, 19.1, 30.9, 31.2, 31.4, 32.1, 68.3, 68.8, 71.5, 100.6, 112.8, 114.3, 114.5, 117.8, 122.1, 123.6, 126.4, 139.6, 148.6, 151.4, 154.7, 156.8, 163.5, 173.4 **(Fig. S2b)**; IR (ν_max_ cm^-1^, ATR): 2954-2870, 1633, 1620, 1596, 1513, 1428, 1325, 1265; ESI-HRMS *m/z* [M+1]^+^ calcd for C_31_H_43_O_6_^+^ 511.3060, found 511.3077.

### Chemistry

The nuclear magnetic resonance (NMR) spectra were measured using the Bruker Avance 600 spectrometer (Bruker, Billerica, MA, USA). The liquid sample was prepared by dissolving each sample (10 mg) in acetone-d6 (0.6 mL). Chemical shifts (δ) is reported in ppm and J values in Hz. Multiplicity is designated as doublet (d), doublet of doublet (dd), triplet (t), and multiplet (m). High-resolution mass spectral analysis (HR-MS) was recorded using the Bruker Impact II instrument (Bruker, Billerica, MA, USA). Calculation of the theoretical molecular mass of compounds was performed using “The Exact Mass Calculator, Single Isotope Version” (http://www.sisweb.com/referenc/tools/exactmass.htm; (Ringoes, NJ, USA)). Infrared spectra (IR) were performed using the IRSpirit-X (Shim-pol) spectrophotometer equipped with the attenuated total reflection (ATR) mode. Melting points were measured by the Electro-thermal IA 9300 melting point apparatus. All commercial substances were purchased in Merck (Darmstadt, German).

### Antioxidant analysis

Fisetin and its derivatives were dissolved in DMSO (1 mg/mL; Merck, Darmstad, Germany). Concentrations in the range of 19.5-500 μM were used for the antioxidant assay. As a reference substance was used the ascorbic acid at the same concentrations such as flavonoids. The antioxidant assay was determined using the previously described method [20, 21]. Briefly, the methanolic solution of 2,2-Diphenyl-1-picrylhydrazyl (DPPH; 100 μL, 3mM) was added to a 96-well plate (Nunc Thermo Fisher Scientific, Waltham, MA, USA). Then, 100 μL of the compound was added to each well and mixed at 30 seconds. After 30 min at 25°C, the absorption wavelength of 515 nm was measured on the BioTek 800TS microplate reader (BioKom, Poland). For all compounds, the activity was carried out at least in triplicate. The values are expressed as the percentages of radical inhibition absorbance (I%) in relation to the control values, as calculated by the following equation: I% = [(A_0_-A_S_/A_0_)x100]

A_0_ is the absorbance of control which exclude the test compounds, and A_S_ is the absorbance of the tested compounds.

### Cell cultures

Cell cultures were performed under standard conditions, at 37 °C in a humidified atmosphere with 5% CO2. All cell lines – CAL-27 (tongue squamous cell carcinoma, ATCC: CRL-2095), A-253 (salivary gland carcinoma; ATCC: HTB-41), UCSF-OT-1109 (tongue squamous cell carcinoma; <u>in the manuscript abbreviated form (UCSF-OT) is used</u>; ATCC: CRL-3442), FaDu (pharynx squamous cell carcinoma; ATCC: HTB-43) and HFF-1 (normal foreskin fibroblast; ATCC: SCRC-1041) come from American Type Culture Collection (ATCC, Manassas, VA, USA). The cells were cultured according to the manufacturer’s instructions, with 1% penicillin-streptomycin solution (Corning Life Science, Painted Post, NY, USA) and 10% fetal bovine serum (FBS; EurX, Poland), except for HFF-1 cell line where 15% FBS concentration was required. All of the culture media (DMEM for CAL-27, UCSF-OT-119 and HFF-1, McCoy’s 5A for A-253 and MEM for FaDu) were obtained from Corning Incorporated (Corning, NY, USA). Cells were routinely cultured in T-25 or T-75 flasks (Fisher Scientific, Thermo Fisher Scientific, Waltham, MA, USA) and then seeded onto 12-well plates (Labsolute, Th. Geyer GmbH, Germany) or Ø 10 mm glass coverslips in a 24-well plate (Labsolute, Th. Geyer GmbH, Germany). In the experiments (except MTT assay), cells were incubated with 10 μM of fisetin (PHL8254, Merck, Darmstadt, Germany) and the derivatives: MKT218 and MKT257 for 24 h in culture medium with a half reduction in FBS concentration, unless stated otherwise.

### Cytotoxicity assay (MTT)

Cells (15,000 cells/well) were seeded in a 96-well plate (Labsolute, Th. Geyer GmbH, Germany) and stimulated with fisetin, MKT218, MKT257 and MKT261 for 24 h at 37°C. After stimulation, cells were incubated with 5 mg/mL MTT reagent (3-(4,5-dimethylthiazol-2-yl)-2,5-diphenyltetrazolium bromide; Merck, Darmstadt, Germany) for ∼1 h at 37°C. Then formazan crystals were dissolved with 100 μL of DMSO. Absorbance was measured at a wavelength of 570 nm using a Varioskan LUX microplate reader (Thermo Fisher Scientific, Waltham, MA, USA).

### Annexin V assay

Cells were seeded (50,000 cells/well) onto 4-chamber culture slides with a glass bottom (Falcon, Corning Inc., Corning, NY, USA) prefixed with 1 % Geltrex LDEV-Free Reduced Growth Factor Basement Membrane Matrix (Gibco, Thermo Fisher Scientific, Waltham, MA, USA). After 24 h of incubation, the cells were preincubated with NucSpot Nuclear stain (Biotium, Fremont, CA, USA) for 1 h. Then, the cells were incubated with Apoptotic, Necrotic and Healthy Cells Quantification Kit (Biotium, Fremont, CA, USA) according to the manufacturer’s instructions. Briefly, the medium was removed, and the cells were stained with 100 μl of binding buffer solution with Annexin V and EthD-III added and incubated for 15 min in dark in RT. Later, cells were washed twice with binding buffer and covered with binding buffer for imaging. Images were obtained with Olympus IX microscope (Olympus Corporation, Tokyo, Japan) at 20x and 40x objective.

### Immunoblotting

Cultured cells were detached with trypsin-EDTA solution (Corning Inc., Corning, NY, USA) in preparation for lysates. Cells were then suspended in cold phosphate-buffered saline (PBS; Corning Inc., Corning, NY, USA) and centrifuged at 300 g. Pellets were resuspended in 200 μl of Pierce RIPA buffer (Thermo Fisher Scientific Waltham, MA, USA) with protease inhibitors (complete Protease Inhibitor Cocktail, Merck, Darmstadt, Germany) and incubated for 30 min at 4 °C with agitation. Lysates were clarified by centrifugation at 1,000 g for 5vmin at 4 °C. Protein concentration was determined by a BCA assay kit (Thermo Fisher Scientific Waltham, MA, USA) to ensure the equal loading of each sample (10 μg of proteins). Samples were electrophoretically separated in a Mini-PROTEAN apparatus (Bio-Rad Laboratories, Hercules, CA, USA) in 10 % SDS-PAGE gels, followed by transfer to a nitrocellulose blotting membrane (Bio-Rad Laboratories, Hercules, CA, USA) in transfer buffer with 20 % methanol. The membranes were blocked with 5% milk dissolved in Tris-buffered saline with 0.1 % Tween 20 (TBST) for 5 min and incubated with anti-PARP1 (ab191217, Abcam; 1:1000) and anti-β actin (4970, Cell Signalling; 1:1000) primary antibodies overnight at 4 °C. The membranes were then washed three times with TBST and incubated with anti-mouse-HRP antibody (Invitrogen, Thermo Fisher Scientific, Waltham, MA, USA; 1:5000) diluted in 5% milk in TBST for 1 h at room temperature. After five washes with TBST, the membrane was incubated with horse-radish peroxidase (HRP) substrate (Bio-Rad Laboratories, Hercules, CA, USA) for chemiluminescence detection that was performed on an Invitrogen iBright FL1500 instrument (Thermo Fisher Scientific Waltham, MA, USA).

### Immunofluorescence staining

Cells cultured on glass coverslips covered with 1% Geltrex (as in the timelapses) were fixed with 4 % Pierce methanol-free formaldehyde (Thermo Fisher Scientific, Waltham, MA, USA) at room temperature for 10 min and followed by two washes with PBS. Cells fixed in formaldehyde were subsequently permeabilized in PBS-Tx at room temperature for 10 min and blocked in 10 % normal donkey serum (Sigma-Aldrich, Merck, Darmstadt, Germany) in 0.1 % PBS-Tx at room temperature for 30 min. Then, the cells were incubated overnight, at 4°C in a humid chamber with primary antibodies diluted in 0.1 % PBS-Tx with 1 % donkey serum. Blocking was followed by three PBS washes before incubation with the appropriate secondary antibodies that were diluted 1:400 in 0.1 % PBS-Tx containing 1 % donkey serum at room temperature for one to 3 h. Finally, after washing with PBS and counterstaining the cell nuclei with 0.5 μg/mL Hoechst (Sigma-Aldrich, Merck, Darmstadt, Germany) for 10 min at room temperature, the coverslips were mounted in Fluorescence Mounting Medium (Dako, Agilent Technologies, Santa Clara, CA, USA) and allowed to dry before imaging. Negative controls with the omission of primary antibodies were performed for each protein. Cells were analysed using the Olympus IX83 microscope with 20x and 40x objective. For the immunocytochemical staining, the following primary antibodies were used: anti-p53 (ab131442, Abcam, Cambridge, UK) diluted 1:200, anti-phospho-H2AX (Ser139, JBW301 clone, 05–636, EDM Milipore, Merck, Darmstadt, Germany) diluted 1:200. The secondary antibodies used were AlexaFluor 488 anti-rabbit IgG (A21206) and AlexaFluor 568 anti-mouse IgG (A10037), both diluted 1:400 (Invitrogen, Thermo Fisher Scientific, Waltham, MA, USA).

### Timelapses

Cells were seeded (50,000 cells/well) onto 4-chamber culture slides with a glass bottom (Falcon, Corning Inc., Corning, NY, USA) prefixed with 1 % Geltrex LDEV-Free Reduced Growth Factor Basement Membrane Matrix (Gibco, Thermo Fisher Scientific, Waltham, MA, USA). After 24 h, NucSpot Live 650 Nuclear Stain (Biotium, Fremont, CA, USA) was added to the cells for 30 min at 37 °C. Then timelapses were performed using Olympus IX microscope with a Lumencor Spectra X 647 nm fluorescence LED light source for 24 h (image intervals – 10 min). Automated Olympus software supported by machine learning was used to calculate the percentage of dead cells (florescence mean (Gray Intensity Value) filtered on 5.000 au, what corresponds to the condensed chromatin signal) and proliferation ratio (calculated by division of 2-24h images frame with 0h images frame).

### Cell cycle (flow cytometry)

Cells were seeded onto 12-well culture plates (Labsolute, Th. Geyer GmbH, Germany). After treatment, cells were detached using trypsin–EDTA solution (Corning, NY, USA), collected, and washed with PBS. Subsequently, cells were fixed by the dropwise addition of ice-cold 70% ethanol while gently vortexing and incubated at −20 °C for a minimum of 2 h prior to analysis. After fixation, ethanol was removed by centrifugation, and cells were washed twice with PBS to eliminate residual ethanol. The cell pellets were then resuspended in 0.5 mL of propidium iodide (PI)/RNase staining solution (BD Pharmingen™ PI/RNase Staining Buffer, BD Biosciences, San Jose, CA, USA) and incubated for 15 min at room temperature in the dark. DNA content was analyzed using a BD Accuri™ C6 Plus flow cytometer (BD Biosciences, San Jose, CA, USA). Each experimental condition was analyzed in three independent replicates. For each sample, 10,000 single-cell events (singlets) were collected and used for subsequent cell cycle distribution analysis.

### Molecular docking study

The X-ray structure of tested protein, such as EGFR (9N6G), PRP1 (7KK4), PI3K (1E7U), AKT1 (6HHF), CDK1 (6GU6), CDK2 (3PXY), CDK4 (9CSK), Bcl-2 (6O0K), Bcl-xl (6UVF), Mcl-1 (4ZBF), GST (1GUL), NQO1 (2F1O), STAT3 (6NUQ), was downloaded from the Protein Data Bank (PDB) [https://www.rcsb.org/]. The structure of tested proteins has been modified by removing crystal water, ions and non-essential ligands. The MGL tools 1.5.6 with AutoDock Vina was used to add Kollman charges to the energy-minimized protein. A grid box was defined around the active site of proteins to encompass key catalytic residues. The structures were saved in a PDBQT format for AutoDock calculations [22]. Ligand structures were generated using ChemSketch and energy-minimized with the MM2 force field [23]. All obtained results were visualized using the BIOVIA Discovery Studio software package [Dessault Systemes BIOVIA. https://www.3dsbiovia.com/products/collaborative-science/biovia-discovery-studio/].

### Statistical analysis

All experiments were performed in duplicate or in triplicate and were repeated independently at least three times, unless otherwise indicated. Data were analyzed with GraphPad Prism 8.0 software. Tests used in the statistical analysis are listed in the description of the figures. Bar graphs represent mean +SEM; ns – non significant, *-p≤0.05, **-p≤0.01, ***-p≤0.001.

## 3. Results

### Increased cytotoxity of alkoxylated fisetin derivatives

In our previous study [10], we demonstrated the anticancer activity of fisetin in head and neck squamous cell carcinoma (HNSCC) cell models. In the present work, we sought to determine whether targeted structural modification of the fisetin molecule could enhance its cytotoxic potential while preserving selectivity toward malignant cells. To this end, we synthesized a series of fisetin analogues bearing substituents containing four carbon atoms, designed to modulate the physicochemical and biological properties of the parent compound. The synthesis was carried out via alkylation of fisetin with appropriate organic bromides in the presence of potassium carbonate in acetone, yielding compounds with four substituted hydroxyl groups (**Fig. 1a**). This approach resulted in the generation of three derivatives: MKT218, containing propynyloxy substituents; MKT257, containing 2-butynyloxy substituents; and MKT261, containing n-butyloxy substituents. The chemical structures of the synthesized compounds were confirmed using a combination of spectroscopic techniques, including proton and carbon nuclear magnetic resonance (^1^H and ^13^C NMR) spectroscopy, infrared (IR) spectroscopy, and high-resolution mass spectrometry (HR-MS).

**Fig. 1.**
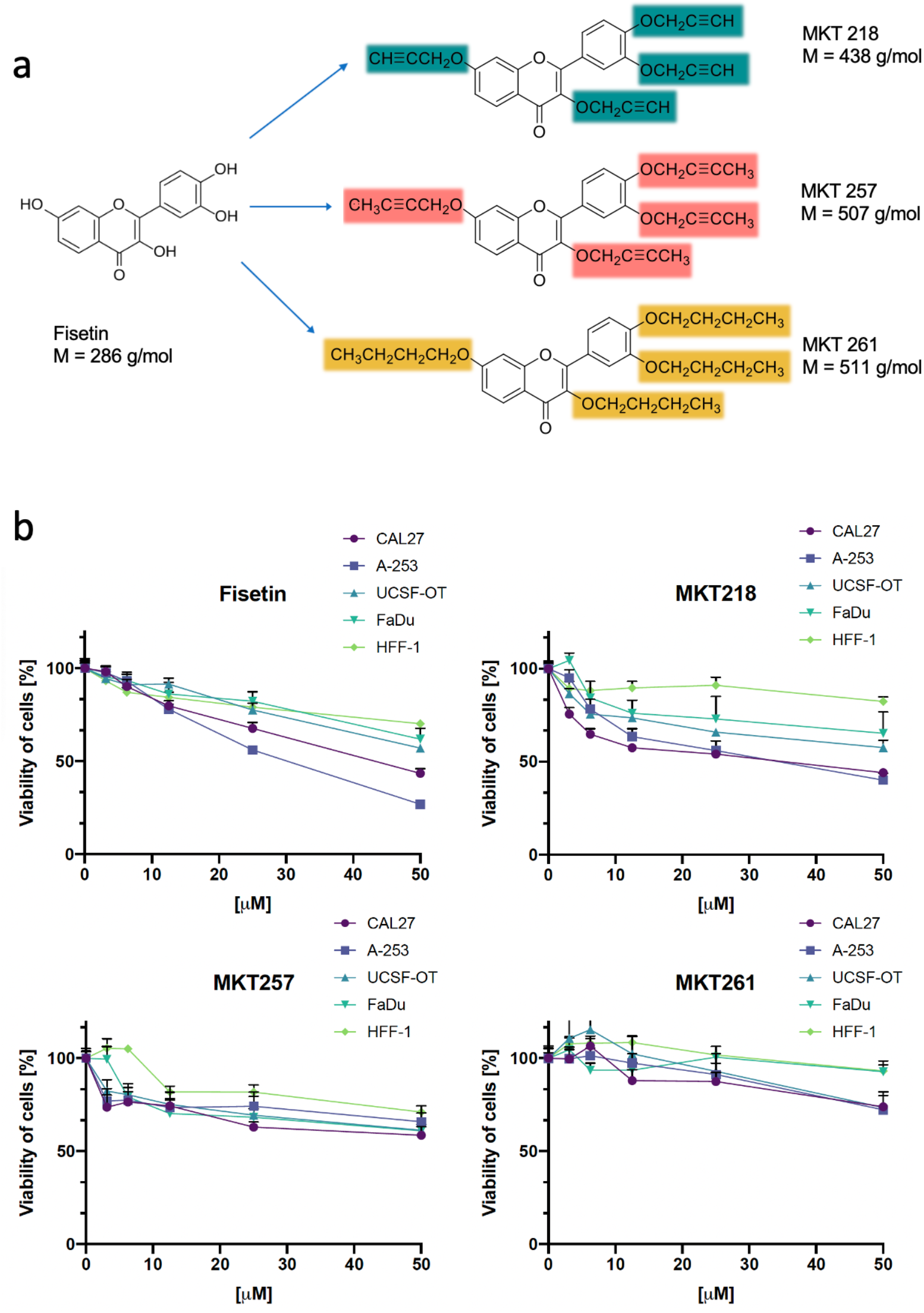
**a)** Schematic representation of fisetin alkoxylated modifications: MKT218, MKT257, MKT 261. **b)** Effect of fisetin and alkoxylated derivatives (3.125-50 μM, 24 h) on cell viability. MTT reduction assay in CAL27, A-253, UCSF-OT, FaDu (cancer cells) and HFF-1 (normal cells). ANOVA. <u>Detailed statistical analysis in supplement</u> (**Fig. S3**).

To assess the cytotoxic potential of the synthesized fisetin derivatives, we performed MTT viability assays across a range of concentrations (3.15, 6.25, 12.5, 25, and 50 μM). Experiments were conducted on four HNSCC cell lines (A-253, CAL27, UCSF-OT, and FaDu), with normal human foreskin fibroblasts (HFF-1) included as a non-malignant control. Quantitative analysis revealed a pronounced and statistically significant reduction in the viability of all tested HNSCC cell lines following treatment with MKT218 at concentrations ranging from 3.15 to 12.5 μM, compared with HFF-1 fibroblasts (**Fig. 1b**). The strongest cytotoxic effects were observed in CAL27 and A-253 cells, in which treatment with 12.5 μM MKT218 reduced cell viability to approximately 60%. Interestingly, within the higher concentration range of MKT218 (12.5–50 μM), a plateau effect was observed in the dose–response curves across all cell types. Despite increased compound concentrations, cytotoxicity toward HFF-1 fibroblasts remained minimal, with cell viability maintained at approximately 90%. The second derivative, MKT257, also induced death of HNSCC cells compared with normal fibroblasts, although this effect was primarily evident at lower concentrations (3.15–12.5 μM) (**Fig. 1b**). At higher concentrations, MKT257 reduced viability in both cancerous and non-cancerous cells. However, the effect remained more pronounced in HNSCC cells, where viability ranged from 58% to 74%, compared with 71% to 81% in HFF-1 fibroblasts. In contrast, the third derivative, MKT261, exhibited substantially weaker cytotoxic activity, even relative to unmodified fisetin. For instance, treatment with 25 μM MKT261 resulted in cell viability ranging between 85% and 100%, irrespective of whether the cells were of malignant or non-malignant origin, indicating a lack of selective anticancer efficacy.

In summary, the fisetin derivatives MKT218 and MKT257 demonstrated significantly enhanced anticancer activity at low concentrations compared with the parent compound fisetin. Based on these findings, a concentration of 10 μM was selected for subsequent experiments, as it provided the strongest cytotoxic effect in HNSCC cells while maintaining low toxicity toward normal fibroblasts. Due to its limited biological activity, the MKT261 derivative was excluded from further analyses. Notably, under the experimental conditions applied in this study, fisetin at a concentration of 10 μM exhibited comparable toxicity toward both normal and cancer cells. This underscoring the improved selectivity of the MKT218 and MKT257 derivatives.

### Enhanced apoptosis activation by alkoxylated fisetin derivatives

To evaluate whether the observed reduction in cell viability is directly associated with enhanced activation of apoptosis induced by fisetin derivatives, we first assessed the cleavage of PARP1, a marker of caspase-dependent apoptotic signalling (**Fig. 2a**). A-253, CAL27, UCSF-OT, and FaDu cell lines were treated with 10 μM fisetin, MKT218, or MKT257 for 24 hours, followed by protein analysis using western blotting. As anticipated, PARP1 cleavage resulting in the characteristic 89 kDa fragment was detected after stimulation with both fisetin and its derivatives. However, no pronounced or statistically significant differences in PARP1 cleavage intensity were observed among the different compounds or across the tested cell lines (**Fig. 2a**). Nevertheless, we did not expect unequivocal results. Western blotting has limited sensitivity of detecting relatively small differences in apoptosis induction, particularly when changes are subtle and occur in only a fraction of the cell population. Rather subtle, possibly a few percent, differences in apoptosis activation required a more sensitive and quantitative method. Therefore, we employed a live-cell microscopy assay utilizing annexin V as a marker of early apoptosis and EthD-III as an indicator of loss of membrane integrity associated with necrosis (**Fig. 2b**). To ensure accurate, artifact-free quantification of correctly stained cells, image analysis was performed using Olympus software equipped with an artificial intelligence-based module. This approach enabled automated segmentation of intact cells and cell nuclei through neural network guided mask generation, allowing precise identification and counting of apoptotic and necrotic cell populations (**Fig. 2b**). We detected a significantly increased proportion of annexin V positive HNSCC cells following treatment with MKT218 (10.7–20.2%) and MKT257 (8.7–15.4%) compared with fisetin-treated counterparts (3.8–9.7%) (**Fig. 2c**). Among the tested cell lines, UCSF-OT cells exhibited the highest sensitivity to MKT218-induced apoptosis, whereas CAL27 cells showed the lowest responsiveness to stimulation. The apoptotic effect induced by MKT257 was only marginally higher than that observed for fisetin. However, its activity appeared more consistent across different HNSCC cell lines. In contrast, analysis of necrotic cell death revealed substantial heterogeneity in cellular responses to fisetin and its derivatives (**Fig. 2c**). UCSF-OT cells did not display a measurable necrotic response following treatment, whereas FaDu cells demonstrated a relatively high proportion of necrotic cells, nearly comparable to the fraction undergoing apoptosis. A-253 and CAL27 cells were more responsive to MKT218 and MKT257, as evidenced by a significantly increased percentage of necrotic cells following stimulation.

**Fig. 2.**
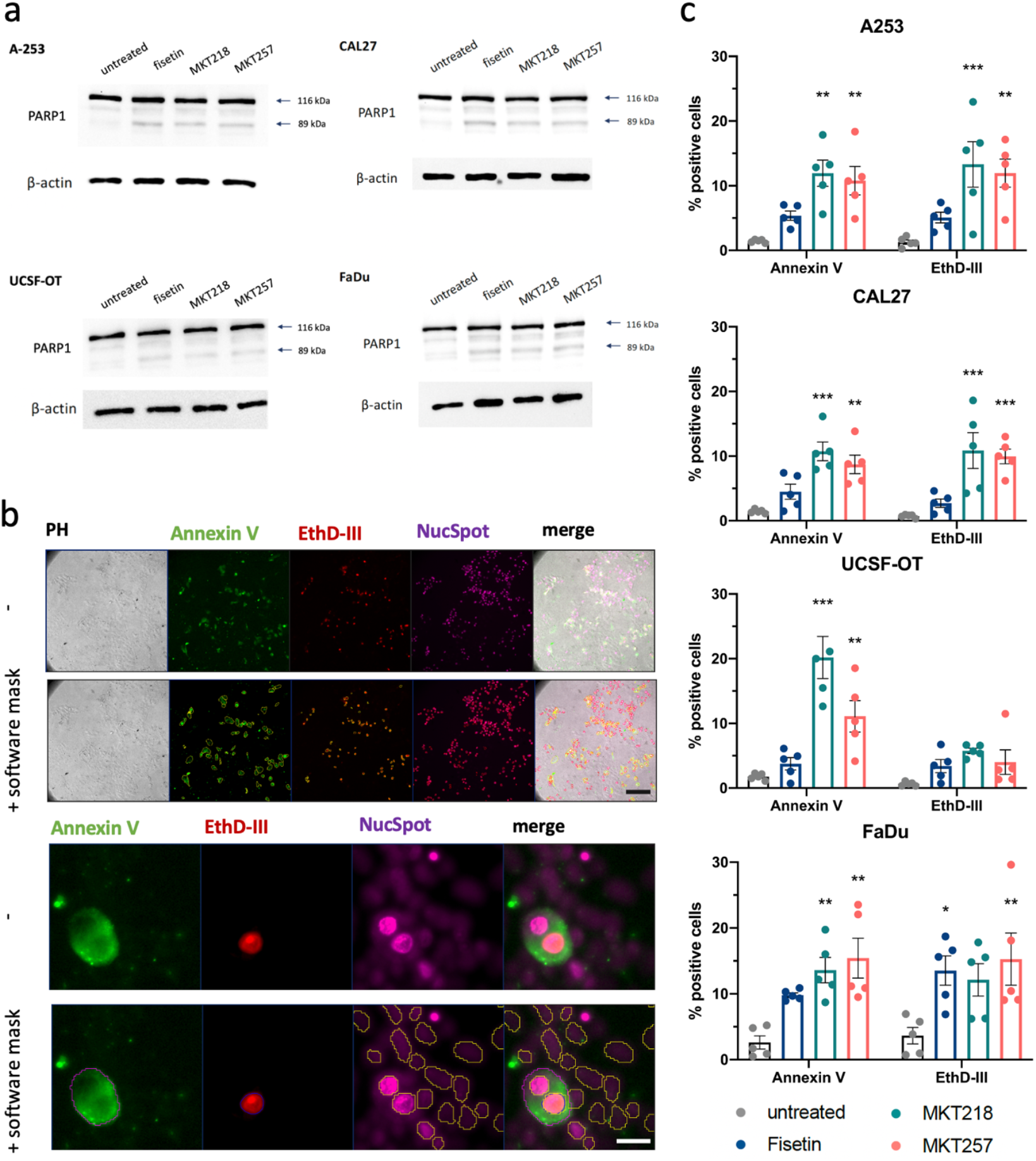
**a)** PARP-1 cleveage in cells treated with fisetin, MKT218 and MKT257 (10 μM, 24 h). Western blotting of PARP1 in A-253, CAL27, UCSF-OT and FaDu cancer cells (cleaved PARP1 was marked as 89 kDa band). β-actin was used as a loading control. **b)** Representative images of Annexin V (green) and EthD-III (red) staining in A-253 cells. 20x magnification images (lower panel digitally zoomed) was analysed by using AI Olympus software – seen as a virtual mask marked on specific cell compartments. Nuclei were counterstained with NucSpot 650 (purple), where the phase contrast (PH) images show the cell morphologies. Fluorescence microscopy. Scale bar: 50 μm (upper panel) and 20 μm (lower panel). **c)** The effect of fisetin, MKT218 and MKT257 (10 μM, 24 h) on apoptosis activation by Annexin V and EthD-III staining in A-253, CAL27, UCSF-OT and FaDu cancer cells. ANOVA. * - untreated vs. treated.

In summary, both fisetin derivatives MKT218 and MKT257, exhibit a markedly higher pro-apoptotic potential compared with the parent compound fisetin. Importantly, as shown in Fig. 1B, this enhanced anticancer activity is accompanied by low cytotoxicity toward normal cells. This highlights the favourable therapeutic profile of these derivatives and supporting their potential as selective anticancer agents.

### Fisetin derivatives do not increase DNA damage response and p53 protein level

Next, we checked whether the observed reduction in HNSCC cell viability and the concomitant increase in apoptotic cell death following treatment with MKT218 and MKT257 were associated with enhanced DNA damage. Thus, we performed immunofluorescence staining (ICC) for phosphorylated histone H2AX (γH2AX), a well-established marker of DNA double-strand breaks (DSBs). In parallel, we assessed the expression level of the tumor suppressor protein p53, which plays a crucial role in the cellular response to genotoxic stress (**Fig. 3a, Fig. S4**). For this purpose, AI-based Olympus image analysis software was employed to enable precise cell segmentation and quantitative evaluation based on p53 and γH2AX fluorescence signal intensities. Quantitative analysis of A-253, CAL27, UCSF-OT, and FaDu cell lines treated with fisetin and its derivatives revealed no statistically significant increase in p53 protein levels when compared with untreated control cells (**Fig. 3b**). Notably, in UCSF-OT and A-253 cells exposed to fisetin, a decrease in p53 expression was observed, which may be attributable to enhanced proteasomal degradation of the protein. This observation is in agreement with previously published reports by other research groups [24]. Similar reduction in p53 signal was observed in all tested HNSCC treated with MKT218 and MKT257. Importantly, no statistically significant increase in γH2AX levels were detected following treatment with fisetin, MKT218, or MKT257 (**Fig. 3b**), indicating the absence of elevated DSB formation.

**Fig. 3.**
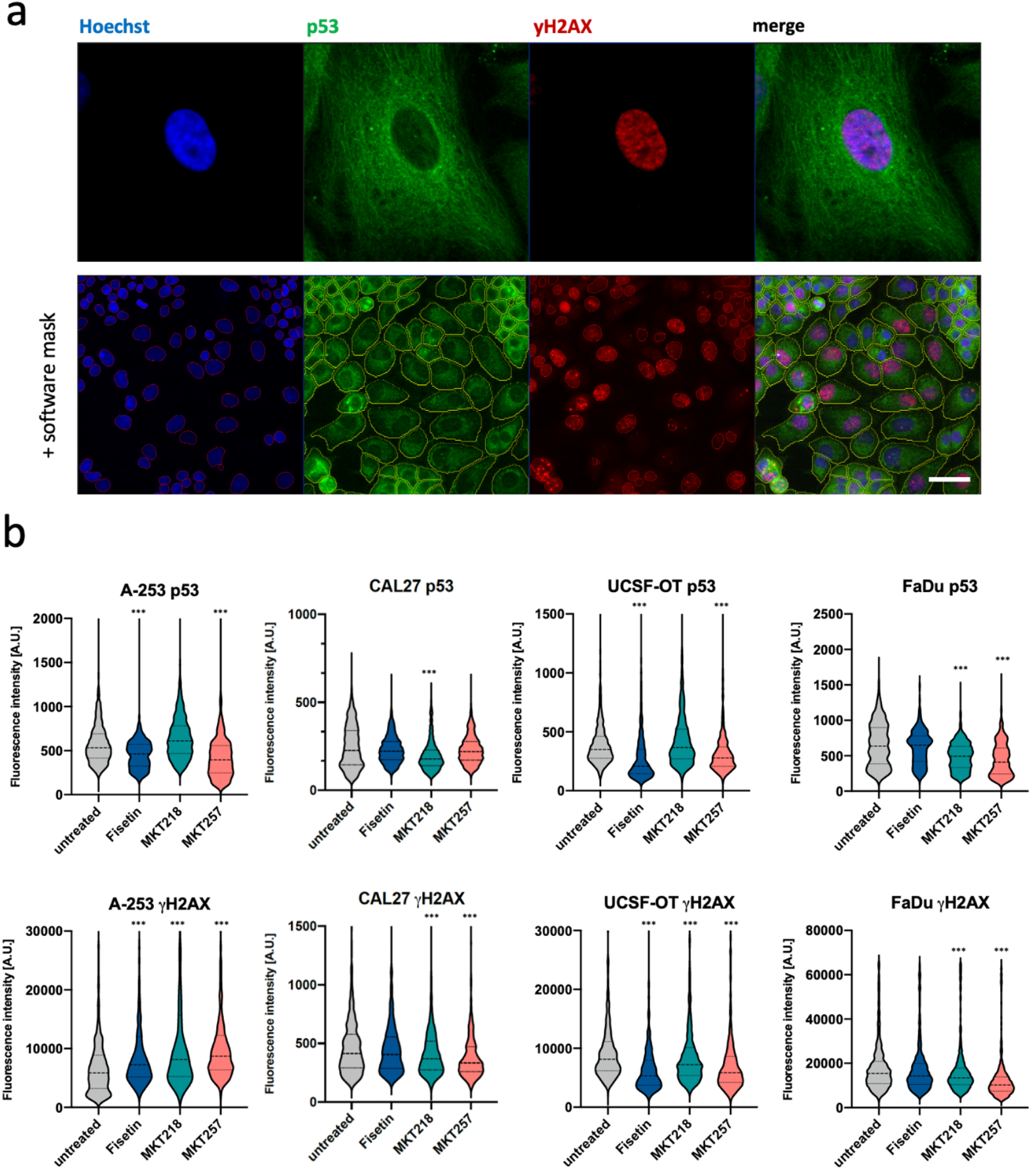
p53 level and DNA-damage response (DDR) in HNSCC. **a)** Representative images of p53 (green) and γH2AX (red) staining in UCSF-OT cells. Cell nucei were counterstained with Hoechst (blue). Zoomed cell (upper panel) and 40x magnification (lower panel) images was analysed by using AI Olympus software – seen as a vitrual mask marked on specific cell compartments. Scale bar: 20 μm. **b)** p53 and γH2AX (DDR marker) staining in A-253, CAL27, UCSF-OT and FaDu exposed to fisetin, MKT218, MKT257 (10 μM) for 24 h. Immunocytochemistry and fluorescence microscopy. Kruskal-Wallis test. * - untreated vs. treated

To summarize, our findings demonstrate that the previously observed reduction in HNSCC cell viability and induction of apoptosis are not mediated by DNA damage dependent mechanisms. Consequently, fisetin and its derivatives are likely to exert their anticancer effects through alternative cellular pathways that selectively target cancer cells rather than by inducing genotoxic stress.

### The anti-proliferative effect of fisetin derivatives

Since the reduced viability of HNSCC cells was not associated with activation of DDR, we next investigated whether treatment with fisetin derivatives affects the proliferative capacity and cell cycle progression. We performed time-lapse microscopy analysis of DNA-stained cells using the NucSpot 650 dye (**Fig. 4a,b**). This experimental approach enabled continuous monitoring of individual cells at 10-minute intervals over a 24 h period. Thus, cell fate and division dynamics after stimulation with fisetin or its derivative might be precisely tracking. Time-lapse analysis revealed a marked increase in the proportion of cells that entered mitosis and successfully formed a metaphase plate but subsequently failed to complete the cell cycle. Instead of progressing to anaphase and cytokinesis, these cells underwent morphological changes characteristic of apoptosis. Such cellular shrinkage and chromatin condensation was accompanied by an increase in nuclear fluorescence intensity (**Fig. 4a**). These observations suggest that fisetin derivatives might induce apoptotic cell death during or immediately following mitotic arrest. Quantitative analysis of time-lapse recordings, performed using AI-assisted Olympus image analysis software, demonstrated a significantly higher percentage of dead cells over the 24 h observation period in all HNSCC cell lines treated with fisetin derivatives compared with untreated controls (**Fig. 4a**). The most pronounced effect was observed following treatment with MKT218, with the percentage of dead cells after 24 hours reaching 32.8% in A-253 cells, 20.6% in CAL27 cells, 19.7% in FaDu cells, and 43.7% in UCSF-OT cells. In contrast, treatment with MKT257 resulted in a stronger cytotoxic effect compared with fisetin only in CAL27 (15.0%) and FaDu (16.8%) cell lines. For reference, under control conditions, the proportion of dead cells ranged from 3.3% to 9.6% across the tested HNSCC models.

**Fig. 4.**
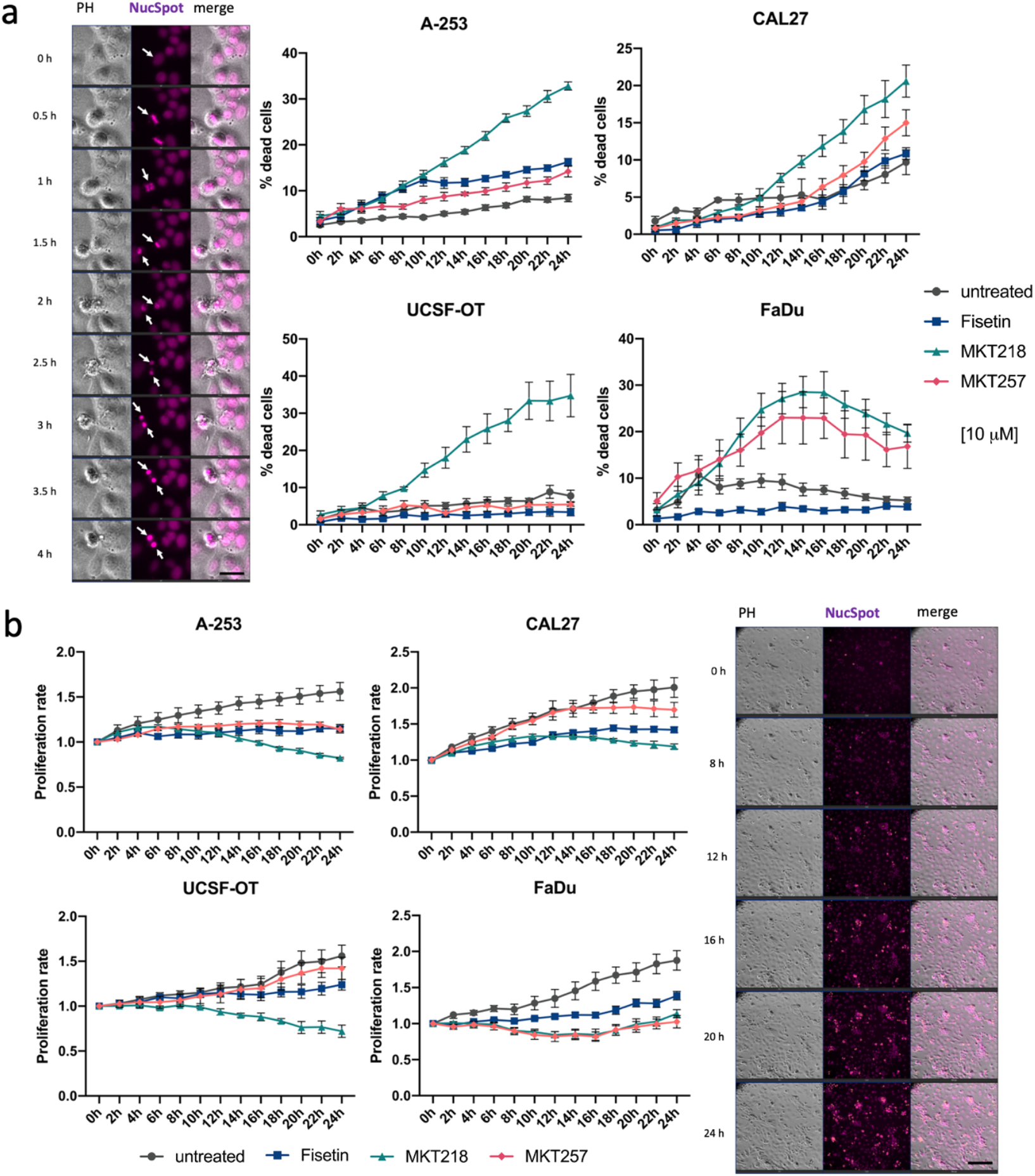
Dead cells level and proliferation rate measured with timelapses of A-253, CAL27, UCSF-OT and FaDu treated with fisetin, MKT218 and MKT257 (10 μM). **a)** Representative images of A-253 cells during 4 h timelapse (left panel). White arrow demonstrates disturbed cell division after formation of the metaphase plate and increased chromatin condensation, which confirm cell death. Nuclei were stained with NucSpot 650 (purple), where the phase contrast (PH) images show the cell morphologies. Objective magnification 20x; digitally zoomed. Scale bar: 20 μm. Increased level of dead cells after MKT218 treatment is observed in all used cancer cells (right panel) comparing to fisetin counterparts. ANOVA **b)** Reduced proliferation rate in all HNSCC treated with MKT218 (left panel). This corresponds to the increased number of cells with condensed chromatin (right panel). In contrast, nearly complete inhibition of HNSCC proliferation after fisetin treatment was observed. ANOVA. Objective magnification 20x. Scale bar: 50 μm. <u>Detailed statistical analysis in the supplemen</u>t (**Fig. S5 - S8**).

In parallel, we evaluated the proliferative potential of HNSCC cells under the same experimental conditions by quantifying changes in cell number over time (**Fig. 4a**). As expected, treatment with MKT218 and MKT257 led to a significant reduction in proliferation rates, which correlated with the increased incidence of apoptosis observed in the time-lapse analysis. Notably, stimulation with fisetin alone resulted in an almost complete flattening of the proliferation curves, suggesting a predominantly cytostatic effect rather than direct induction of cell death. This observation implies that fisetin may primarily interfere with cell cycle progression, thereby limiting proliferation without strongly triggering apoptosis. Among the tested compounds, MKT218 exerted the most pronounced inhibitory effect on cell proliferation, with the greatest reduction in proliferation rate observed in A-253 (0.82) and UCSF-OT (0.72) cell lines (**Fig. 4b**). These findings further support the superior anticancer activity of this derivative compared with both fisetin and MKT257.

In summary, fisetin derivatives significantly promote apoptotic cell death in HNSCC cells with simultaneous suppression of cellular proliferation. These effects are substantially more pronounced comparing to the parent compound fisetin, which, in our experimental setup, appears to act primarily through inhibition of cell proliferation rather than induction of apoptosis.

### Fisetin derivatives do not affect cell cycle

The disturbance in the proliferation rate of HNSCC cells stimulated with fisetin and its derivatives suggests abnormalities in G1/S/G2 phases of the cell cycle. Hence, we performed cytometric analysis of the cell cycle using propidium iodide (**Fig. 5a**). The most pronounced biological effect was observed following fisetin treatment in the CAL-27, UCSF-OT, and FaDu cell lines. Fisetin induced a marked cell cycle arrest at the G2/M phase compared to control cells. In the CAL27 cell line, the proportion of cells in the G2/M phase increased from 19.6% in control conditions to 34.7% following fisetin treatment (**Fig. 5b**). A similar effect was observed in the UCSF-OT cell line, where the G2/M population increased from 17.78% to 29.17%, as well as in the FaDu cell line, where it increased from 19.13% to 34.13%. Additionally, in the FaDu cell line, fisetin treatment resulted in an increased proportion of cells in the S phase of the cell cycle, rising from 13.67% in control cells to 21.72% in treated samples, suggesting a possible impairment of cell cycle progression at the DNA synthesis stage (**Fig. 5b**). In these three cell lines, the fisetin derivatives MKT218 and MKT257 induced only subtle alterations in cell cycle distribution, without a clear accumulation of cells in any specific phase. A distinct response profile was observed in the A253 cell line. In this case, the most pronounced effect was detected following treatment with the MKT218 derivative, which led to an increase in the proportion of cells in the G2/M phase from 24.26% in control cells to 32.56% (**Fig. 5b**). This represented the most substantial change observed in this cell line, while the remaining compounds induced only minor shifts in cell cycle phase distribution. Analysis of the subG1 population, indicative of DNA fragmentation and potential apoptotic processes, demonstrated that fisetin exerted the strongest effect in this fraction. However, the maximal observed percentage of subG1 cells was only 3.29% (in the FaDu cell line).

**Fig. 5.**
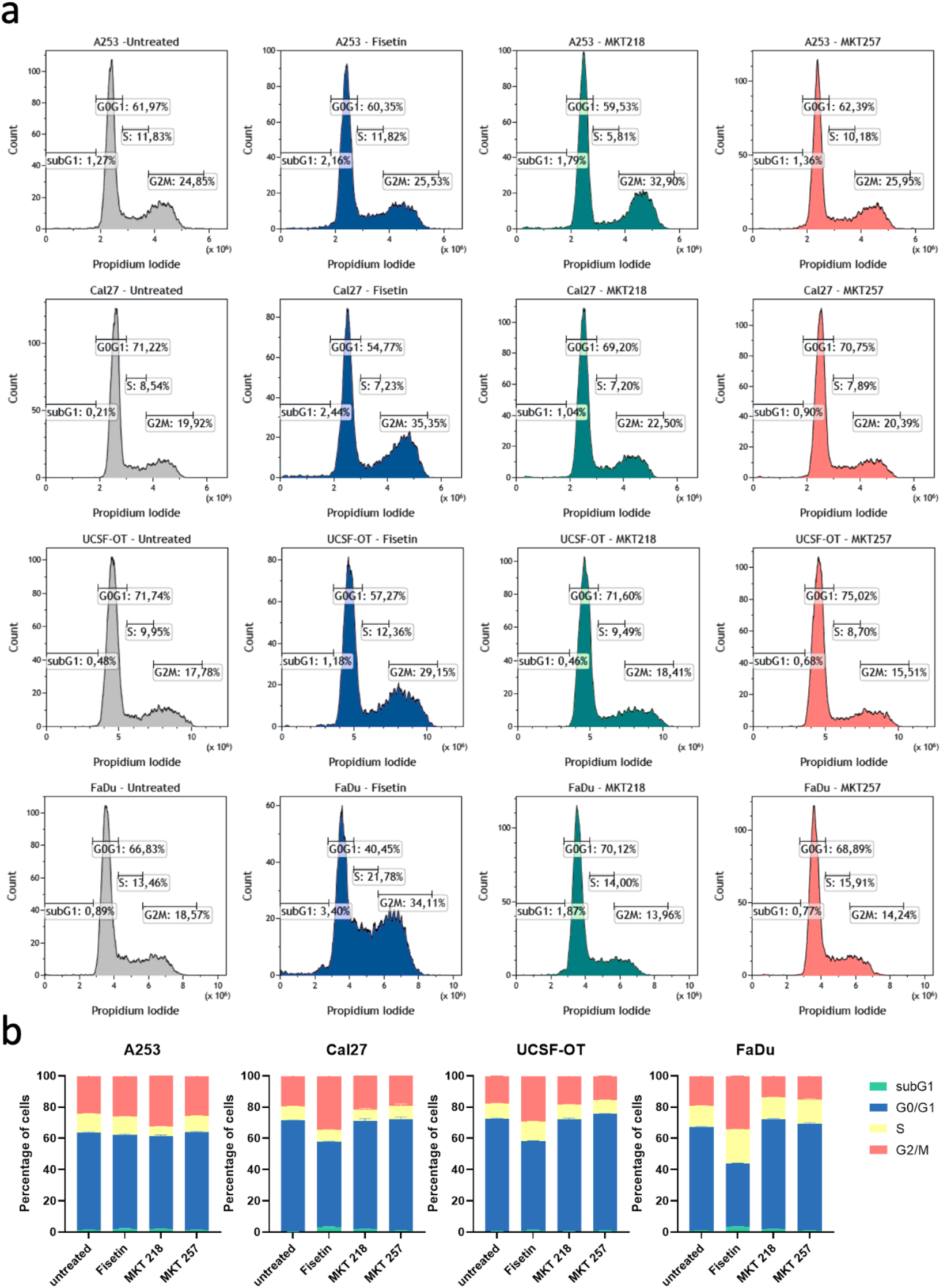
Cell cycle distribution in A-253, CAL27, UCSF-OT, and FaDu cell lines treated with fisetin MKT218 and MKT257 (10 μM) for 24 h. **a)** Representative cell cycle histograms obtained by flow cytometry and (**b)** stacked bar charts summarizing the percentage distribution of cells in the subG1, G0/G1, S, and G2/M phases of the cell cycle.

In conclusion, we confirmed that fisetin primarily inhibits HNSCC cell growth by inducing cell cycle arrest in G2/M phase. In contrast, the fisetin derivatives MKT218 and MKT257 generally did not affect HNSCC cell cycle phases, what may suggest their mechanisms of action differ from the parent compound. Collectively, these results suggest that spatial and structural chemical modifications of fisetin may potentially influence biological activity of different proteins involved in regulation of critical cellular processes.

### Increased molecular docking of fisetin derivatives to anti-apoptotic proteins active sites

All results obtained to date consistently suggest that the fisetin derivatives MKT218 and MKT257 operate through cellular mechanisms that are distinct from those of the parent compound. Consequently, the induction of cell death observed in response to these derivatives may be linked to differences in their spatial orientation and interaction with specific target proteins. In this context, our analysis focused on a set of key proteins involved in the regulation of the cell cycle, including CDK1, CDK2, and CDK4, as well as proteins that play central roles in cell survival and anti-apoptotic signalling pathways, such as PI3K, AKT1, Bcl-2, and Bcl-XL (**Fig. 6a**). We performed molecular docking, which allowed us to verify the binding energy of fisetin and its derivatives to the active sites of proteins. The result was presented as a scoring value (ΔG) which determines the binding energy between protein and ligand (**Fig. 6a**). The lower value of ΔG means that the ligand is better fitted to the active centre of the protein. Comparison of the biological results and computer simulation shows that fisetin creates a more stable complex with cyclin-dependent kinase (CDK) proteins family, which are involved in cell cycle arrest in G2/M phase, than MKT218 and MKT257. Moreover, the obtained ΔG values are similar to this obtained for Abemaciclib. As seen on **Fig. 6b**, fisetin is located deep in the hydrophobic matrix of the protein, while the 1-(pyridine-3-ylmethyl)piperazine fragment of Abemaciclib is localized outside the active centre of protein. This arrangement suggests that complex of fisetin with CDK proteins may be more permanent than Abemaciclib-CDK complex (**Fig. S9, Fig. S10**). On the other hand, introduction of substituents with triple bond increases the value of ΔG toward PI3K, AKT1, Bcl-2, and Bcl-XL. Comparing the arrangement of fisetin in the active site of tested proteins shows that hydroxyl group at C4’ position of fisetin creates a hydrogen bond, while flavone scaffold interacts with amino acid by hydrophobic interaction. In the complex between MKT218 and MKT257 and Bcl-2 protein, the carbonyl group creates a hydrogen bond with ARG146, while flavone scaffold and substituents create hydrophobic interaction with VAL148, VAL133, ALA149, GLY146 and PHE104 (**Fig. S11**). In the active site of Bcl-XL and AKT1 protein the ligand creates only hydrophobic interaction (**Fig. S12 and S13**).

**Fig. 6.**
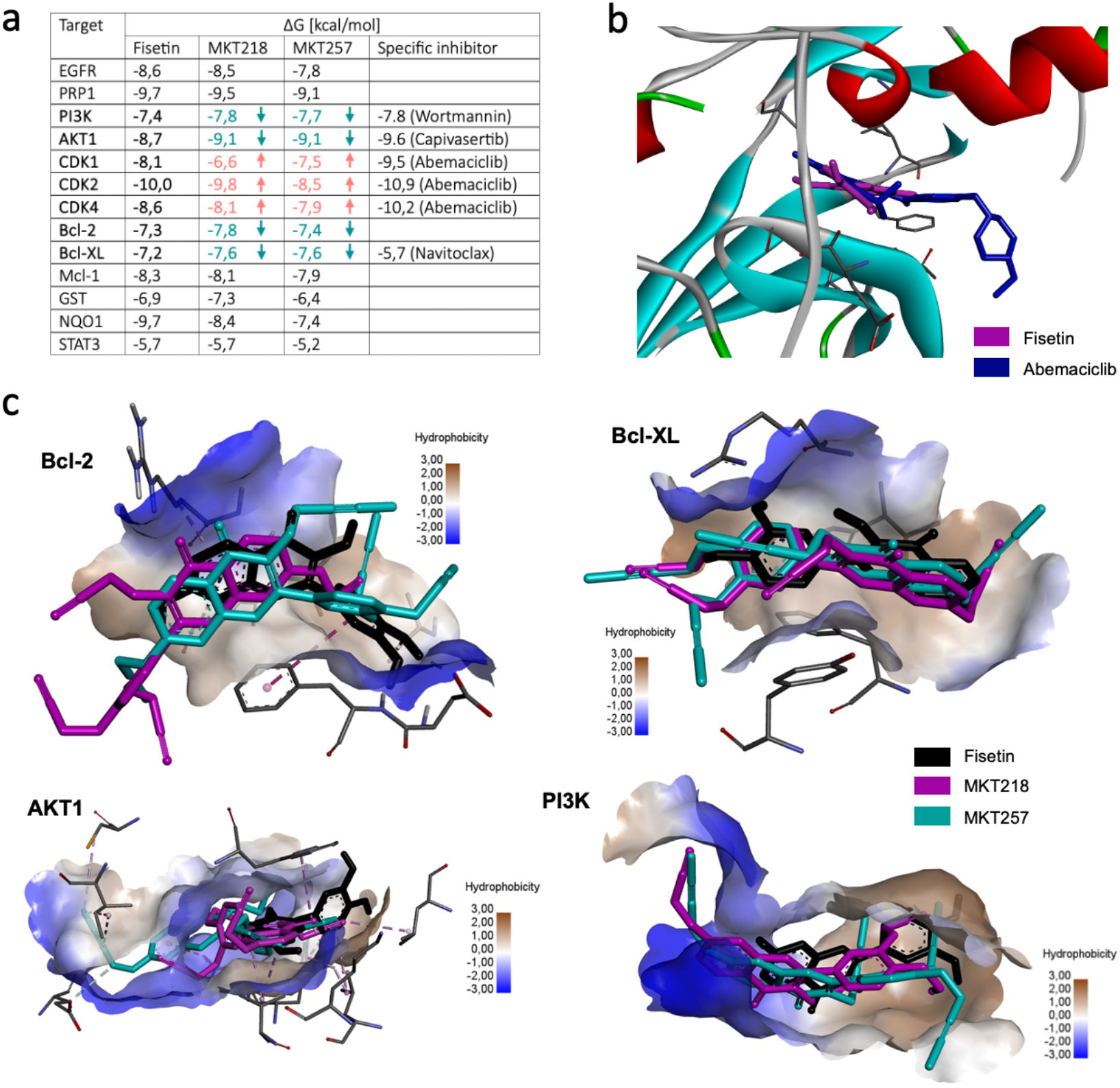
Docking analysis of fisetin, MKT218 and MKT257. **a)** Table representing the binding energy (ΔG) between selected proteins and fisetin, MKT218, MKT257 and selected inhibitors. **b)** Molecular docking comparison between fisetin and Abemaciclib in CDK proteins. **c)** Molecular docking of fisetin, MKT218 and MKT257 with Bcl-2, Bcl-XL, AKT1 and PI3K proteins.

To summarize, our results indicate that MKT218 and MKT257 act via mechanisms distinct from fisetin, potentially due to differences in their spatial conformation and protein interactions. Molecular docking of fisetin showed more stable binding to CDK proteins consistent with its role in G2/M cell cycle arrest. In contrast, the derivatives display increased affinity toward survival-related proteins.

## 4. Discussion

Fisetin is a naturally occurring flavonoid that has attracted considerable interest due to its reported anticancer properties across multiple tumor types, including head and neck squamous cell carcinoma (HNSCC). In our previous work [10], we demonstrated that fisetin reduces the viability of HNSCC cells; however, its limited selectivity toward cancer cells and relatively modest pro-apoptotic activity highlighted the need for further optimization. Other groups reported that fisetin promotes apoptotic cell death via both intrinsic (mitochondrial) and extrinsic (death receptor–mediated) pathways across multiple cancer models, with caspase activation and PARP cleavage consistently reported in *in vitro* systems and xenograft studies [15, 25]. In addition, fisetin induces cell-cycle arrest at the G2/M or G1/S checkpoints and modulates key kinases involved in the control of tumor cell proliferation [15].

In the present study, we investigated whether rational structural modification of fisetin could enhance its anticancer efficacy while preserving low toxicity toward normal cells. Chemical modification of fisetin by introducing four-carbon substituents resulted in derivatives with distinct biological activities. Among the synthesized compounds, MKT218 (propynyloxy substituents) and MKT257 (2-butynyloxy substituents) significantly reduced HNSCC cell viability at low micromolar concentrations, whereas MKT261 (n-butyloxy substituents) showed minimal cytotoxic activity. These observations suggest that not only the length but also the degree of unsaturation and structural rigidity of the substituents critically influence the biological activity of fisetin derivatives. Importantly, both MKT218 and MKT257 demonstrated enhanced selectivity toward cancer cells, as normal human fibroblasts retained high viability at concentrations that were cytotoxic to HNSCC cells. In our previous work MKT218 was used in different experimental set-up with working concentrations tailored to match ½ IC_50_ and ¼ IC_50_ established with MTT assay [18]. Moreover, the cells were previously treated with a culture medium devoid of FBS, and here we used a reduced concentration (5% FBS). This experimental modification allowed us to reduce the proliferative effect of HNSCC cells and, on the other hand, to reduce the uptake of fisetin and fisetin derivatives by albumins [26]. However, the presence of serum may have influenced cellular metabolic activity in the MTT assay (**Fig. 1b**), potentially contributing to an apparent plateau in the viability curve at higher concentrations of the derivatives. Hence the concentration of 10 μM proposed in the experiments seems to be the most appropriate.

The reduced viability of HNSCC cells following treatment with fisetin derivatives was not associated with activation of a canonical DNA damage response (DDR). Neither increased γH2AX levels nor upregulation or nuclear accumulation of p53 was observed across multiple HNSCC cell lines, indicating that double-strand DNA breaks (DSB) do not contribute significantly to the observed cytotoxic effects. In contrast, fisetin-induced DSB has been shown in other HNC cell lines (CAL-33, UM-SCC-22B, HSC-3), working in a time- and concentration-dependent matter, though the concentrations were significantly higher (25, 40, 50 μM) than the ones used in our experiment (10 μM) [13, 27]. Khozooei *et al*. reported increased DSB in breast cancer cells following treatment with high concentrations of fisetin, with this effect being further potentiated by concomitant exposure to 4 Gy ionizing radiation [28]. Notably, these observations were obtained using a substantially higher fisetin concentration (75 μM). In addition, in our experimental setup, a reduced serum content was applied to the culture medium to limit fisetin sequestration by albumin. Shih *et al*. showed that in HSC-3 cells, stimulation with high concentrations of fisetin induced a significant increase in reactive oxygen species (ROS) levels after 48 hours [27]. However, our results using the DPPH assay suggest that fisetin may act as an antioxidant, practically as strongly as ascorbic acid (**Fig. S14**). Moreover, other studies also demonstrate a protective effect of fisetin against oxidative stress, including in breast cancer [29], liver cancer [30], or cervical cancer [31]. Therefore, we did not expect increased oxidative DNA damage. Interestingly, fisetin derivatives MKT218 and MKT257 did not reduce free radicals in the DPPH assay (**Fig. S14**) but still had no effect on DSBs.

The absence of a DDR suggests that fisetin derivatives act through alternative, non-genotoxic mechanisms. Nevertheless, our data indicate that enhanced apoptosis induction is a key contributor to the anticancer activity of fisetin derivatives. Although PARP1 cleavage detected by western blotting confirmed activation of apoptotic signalling. PARP-1 cleavage pattern is consistent with other published data, where a significantly greater cleavage is detected in concentrations of 25 μM and more [32]. However, we did not observe differences between fisetin, MKT218 and MKT257. Theoretically, the reduction in viability of HNSCC cells treated with MKT218 and MKT257 suggested a stronger cleavage of PARP1, hence we used a more sensitive technique. Live-cell imaging combined with annexin V and EthD-III staining, supported by AI-assisted image analysis, revealed a significantly higher proportion of apoptotic cells following treatment with MKT218 and MKT257, compared with fisetin. Notably, MKT218 consistently induced the strongest apoptotic response across all tested HNSCC cell lines, underscoring its superior pro-apoptotic potential.

Subsequently, time-lapse microscopy provided direct observation of cell fate following compound treatment. Fisetin derivatives, particularly MKT218, increased the proportion of cells that entered mitosis but failed to complete cell division (**Fig. 4a**). These cells exhibited hallmark features of apoptosis, including cellular shrinkage and chromatin condensation, suggesting that fisetin derivatives trigger apoptotic cell death during or shortly after mitotic arrest. This phenomenon resembles mitosis-associated apoptosis, a mechanism that selectively affects rapidly proliferating cancer cells and is often exploited by anticancer agents targeting cell cycle progression [33]. In contrast, fisetin itself exerted a predominantly cytostatic effect, as evidenced by the pronounced flattening of proliferation curves in time-lapse experiments, with comparatively limited induction of apoptosis. This observation confirms that fisetin primarily interferes with cell cycle progression rather than actively promoting cell death. Respectively, Smith *et al*. showed that fisetin caused G2/M arrest with reduced histone H3 Ser10 phosphorylation implicating Aurora B kinase inhibition [15]. Our data also shows that fisetin predominantly acts as a cytostatic agent, as evidenced by a pronounced accumulation of cells in the G2/M phase observed in multiple cell lines (**Fig. 5a**). Accordingly, Xianghua *et al*. showed that fisetin downregulated cyclin-dependent kinases (CDK2, CDK4) and shifted phosphorylation state of the retinoblastoma proteins from hyperphosphorylated to hypophosphorylated in colorectal models [34]. In contrast, the fisetin derivatives MKT218 and MKT257 induced only minor alterations in cell cycle phase distribution in most HNSCC models. Despite their strong anti-proliferative effects, these compounds did not cause a clear arrest at G1, S, or G2/M phases, suggesting that they do not primarily target classical cell cycle checkpoints.

Molecular docking did not identify a single protein target for which MKT218 and MKT257 exhibits a markedly higher binding affinity compared to fisetin. However, MKT218 shows consistently comparable or slightly more favourable predicted binding free energies toward several proteins involved in cell survival and apoptosis, including PI3K, AKT1, Bcl-2, and Bcl-XL (**Fig. 6**). Although the observed differences in ΔG values are generally modest and remain within the typical uncertainty of docking methods, their convergence on functionally related targets may be of potential biological relevance. Importantly, PI3K/AKT1 signaling is crucial in anticancer treatment, particularly in the acquisition of chemoresistance, for example, by FGF-1 [35]. Such molecular docking could therefore counteract this effect. In contrast, fisetin displays stronger predicted binding to targets associated with cell cycle regulation and redox processes, such as CDK1 and NQO1. This is consistent with our both propidium iodide analyses and the DPPH assay in HNSCC. Overall, these results suggest that the enhanced cytotoxic activity of MKT218 may be associated with a broader, multi-target interaction profile rather than strong inhibition of a single molecular target, warranting further experimental validation.

Importantly, other research groups for years have explored the modifications of fisetin to improve its efficacy against cancer cells. For instance, Sabarwal *et al*. have synthesised a 4’-brominated derivative of fisetin against non-small-cell lung carcinoma. The compounds have induced apoptosis, G2/M cell cycle arrest and inhibition of proliferation of A549 and H1299 cell lines, although a comparison with fisetin or a healthy cell line was not conducted [36]. Another group by Rzeszutek *et al*. have developed mitochondria-targeted fisetin derivatives. Almost all of the synthesised compound showed a stronger cytotoxic effect when compared with fisetin, but one (mF3) was selected for further examination. The compound has induced apoptosis and cell death of ER-positive breast cancer at concentrations as low as 1 μM, without affecting healthy control epithelial cells [37]. However, research on fisetin derivatives remains limited, resulting in a small number of studies available for result comparison. Interestingly, the mechanism of fisetin activity might me related to its metabolite rather than the original compound. Group led by Touil *et al*. have evaluated the pharmacokinetics of fisetin in mice, and found that a methylated metabolite of fisetin, geraldol, exhibited a stronger deposition in tumour post-injection than fisetin or any found metabolite. Additionally, geraldol had a stronger cytotoxic effect on Lewis lung carcinoma cell lines than fisetin, with concurrent safer cytotoxic profile as per EAhy 926 and NIH 3T3 normal endothelial cells [38]

Taken together, our results demonstrate that targeted alkoxylated modification of fisetin can substantially enhance its anticancer properties by increasing apoptotic potential and improving selectivity toward cancer cells. Among the tested compounds, MKT218 emerged as the most promising derivative, combining strong pro-apoptotic activity, suppression of proliferation, and low toxicity toward normal cells. These findings provide a strong rationale for further investigation of fisetin derivatives as potential therapeutic agents for HNSCC. However, including in-depth mechanistic studies and evaluation in more complex preclinical *in vitro* and *in vivo* models have to be implemented.

## 5. Data availability

The data supporting the conclusions of this article are available from the corresponding author upon reasonable request.

## 6. Abbreviations

HNSCC: head and neck squamous cel carcinoma
HNC: squamous cell carcinoma
HPV: human papilloma virus
DPPH: 2,2-Diphenyl-1-picrylhydrazyl
CDKIs: cyclin-dependent kinase inhibitors
PARP-1: Poly(ADP-Ribose) Polymerase 1
TLC: thin layer chromatography
NMR: nuclear magnetic resonance
IR: infrared spectra
ATR: attenuated total reflection
DMSO: dimethyl sulphoxide
FBS: fetal bovine serum
TBST: Tris-buffered saline with 0.1 % Tween 20
PBS: phosphate-buffered saline
HRP: horseradish peroxidase
PI: propidium iodide
PDB: Protein Data Bank
ICC: immunofluorescence staining
DDR: DNA-damage response
DBSs: double-strand breaks
CDK: cyclin-dependent kinase
ROS: reactive oxygen species

## 8. Funding

This work was supported by Medical University of Silesia grants: BNW-1-030/N/4/F (R.K.) and PCN-1-203/K/1/I (A.D.)

### Contributions

Study design: PC. Methodology: PC. Validation: PC. Experiments: PC, PW, KK. Data analysis: PC. Visualization: PC. Manuscript preparation: PC, PW, KK, MKT. Draft revision: PC, PW, KK, MKT, AD, RK. Supervision: RK. All authors read and approved the final manuscript.

