## Supplemental Figures for "Alkoxylated fisetin derivatives – the new potential drugs against head and neck cancers"

### Supplementary materials

a

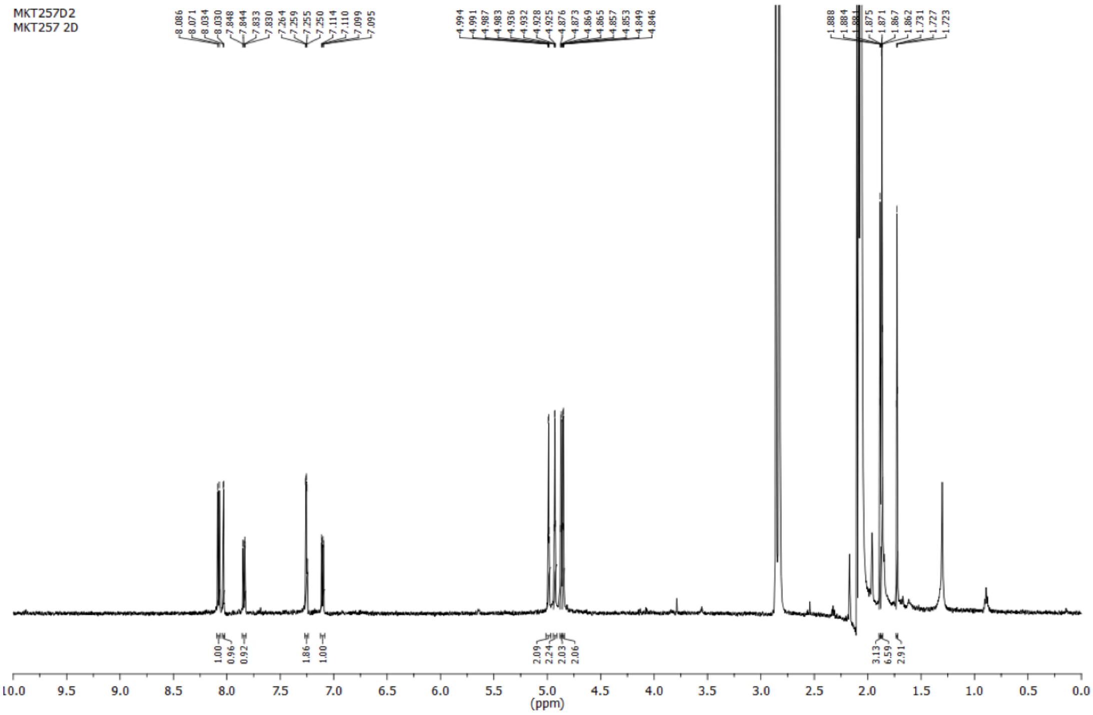

b

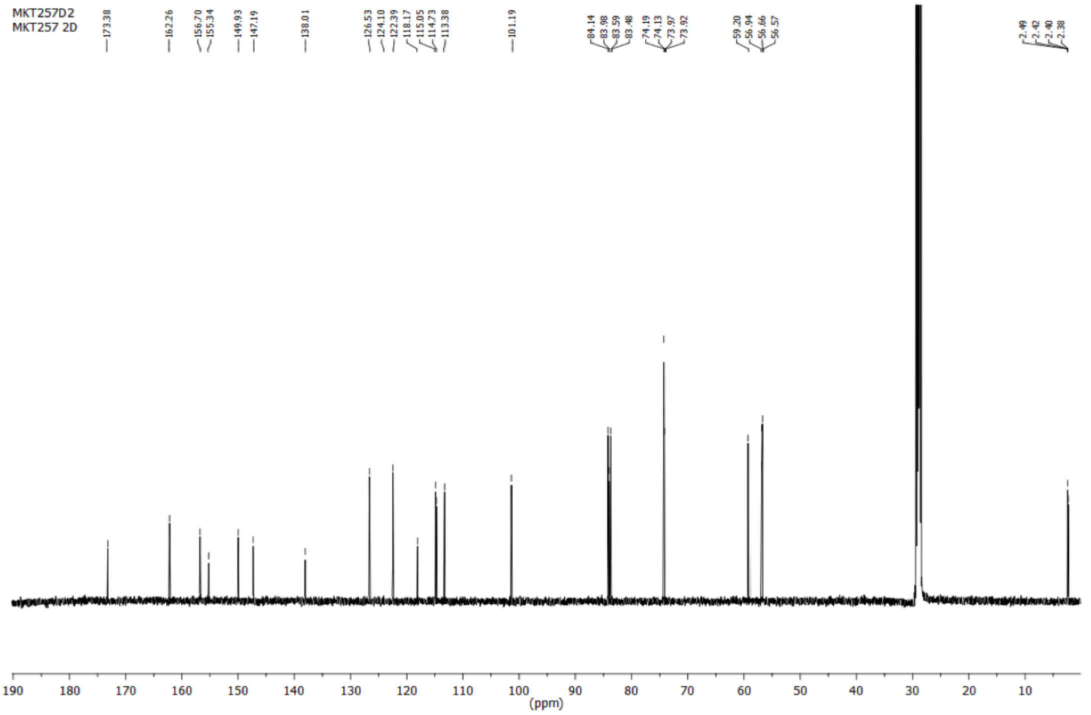

**Fig. S1 a**) $^{1}H$ NMR and (**b**) $^{13}C$ NMR spectra of 2-(3,4-bis(but-2-yn-1-yloxy)phenyl)-3,7-bis(but-2-yn-1-yloxy)-4H-chromen-4-one MKT257.

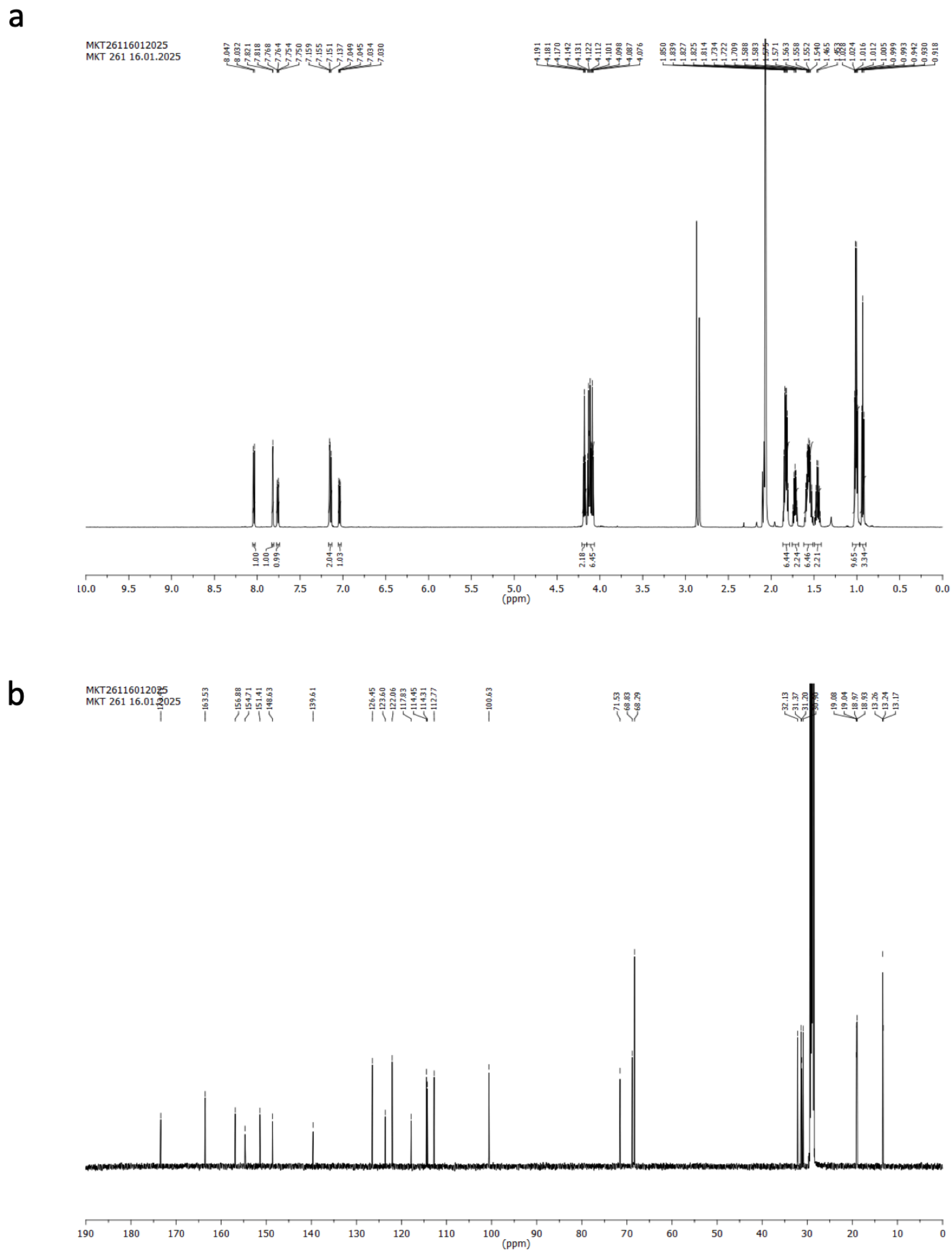

**Fig. S2 a**) $^{1}$H NMR and (**b**) $^{13}$C NMR spectra of 3,7-dibutoxy)-2-(3,4-dibutoxy)phenyl)- 4H-chromen-4-one MKT261.

**a**

| Dunnett's multiple comparisons test | Predicted (LS) mean diff. | 95.00% CI of diff. | Significant? | Summary | Adjusted P Value |
|---|---|---|---|---|---|
| Row 1 | | | | | |
| HFF-1 vs. CAL27 | -1.111e-008 | -12.32 to 12.32 | No | ns | >0.9999 |
| HFF-1 vs. A-253 | 6.656e-006 | -12.32 to 12.32 | No | ns | >0.9999 |
| HFF-1 vs. UCSF-OT | 9.989e-006 | -12.32 to 12.32 | No | ns | >0.9999 |
| HFF-1 vs. FaDu | 9.989e-006 | -12.32 to 12.32 | No | ns | >0.9999 |
| Row 2 | | | | | |
| HFF-1 vs. CAL27 | -4.936 | -17.26 to 7.384 | No | ns | 0.7215 |
| HFF-1 vs. A-253 | -4.340 | -16.66 to 7.980 | No | ns | 0.8003 |
| HFF-1 vs. UCSF-OT | -1.156 | -13.48 to 11.16 | No | ns | 0.9979 |
| HFF-1 vs. FaDu | -2.016 | -14.34 to 10.30 | No | ns | 0.9839 |
| Row 3 | | | | | |
| HFF-1 vs. CAL27 | -3.052 | -15.37 to 9.269 | No | ns | 0.9312 |
| HFF-1 vs. A-253 | -6.000 | -18.32 to 6.320 | No | ns | 0.5705 |
| HFF-1 vs. UCSF-OT | -4.101 | -16.42 to 8.220 | No | ns | 0.8294 |
| HFF-1 vs. FaDu | -6.435 | -18.75 to 5.886 | No | ns | 0.5095 |
| Row 4 | | | | | |
| HFF-1 vs. CAL27 | 4.684 | -7.637 to 17.00 | No | ns | 0.7558 |
| HFF-1 vs. A-253 | 6.388 | -5.932 to 18.71 | No | ns | 0.5160 |
| HFF-1 vs. UCSF-OT | -7.034 | -19.35 to 5.286 | No | ns | 0.4297 |
| HFF-1 vs. FaDu | -1.734 | -14.05 to 10.59 | No | ns | 0.9908 |
| Row 5 | | | | | |
| HFF-1 vs. CAL27 | 11.39 | -1.267 to 24.06 | No | ns | 0.0907 |
| HFF-1 vs. A-253 | 23.01 | 10.35 to 35.67 | Yes | **** | <0.0001 |
| HFF-1 vs. UCSF-OT | 1.471 | -11.19 to 14.13 | No | ns | 0.9954 |
| HFF-1 vs. FaDu | -3.243 | -15.90 to 9.419 | No | ns | 0.9198 |
| Row 6 | | | | | |
| HFF-1 vs. CAL27 | 26.71 | 14.39 to 39.03 | Yes | **** | <0.0001 |
| HFF-1 vs. A-253 | 43.28 | 30.96 to 55.60 | Yes | **** | <0.0001 |
| HFF-1 vs. UCSF-OT | 13.07 | 0.7456 to 25.39 | Yes | * | 0.0338 |
| HFF-1 vs. FaDu | 8.250 | -4.070 to 20.57 | No | ns | 0.2895 |

**b**

| Dunnett's multiple comparisons test | Predicted (LS) mean diff. | 95.00% CI of diff. | Significant? | Summary | Adjusted P Value |
|---|---|---|---|---|---|
| Row 1 | | | | | |
| HFF-1 vs. CAL27 | 1.111e-006 | -14.25 to 14.25 | No | ns | >0.9999 |
| HFF-1 vs. A-253 | -1.111e-006 | -14.25 to 14.25 | No | ns | >0.9999 |
| HFF-1 vs. UCSF-OT | 1.667e-006 | -15.93 to 15.93 | No | ns | >0.9999 |
| HFF-1 vs. FaDu | 1.667e-006 | -15.93 to 15.93 | No | ns | >0.9999 |
| Row 2 | | | | | |
| HFF-1 vs. CAL27 | 13.71 | -0.5331 to 27.96 | No | ns | 0.0632 |
| HFF-1 vs. A-253 | -5.792 | -20.04 to 8.453 | No | ns | 0.7219 |
| HFF-1 vs. UCSF-OT | 2.633 | -13.29 to 18.56 | No | ns | 0.9845 |
| HFF-1 vs. FaDu | -15.23 | -31.15 to 0.6995 | No | ns | 0.0658 |
| Row 3 | | | | | |
| HFF-1 vs. CAL27 | 23.44 | 8.295 to 38.58 | Yes | *** | 0.0008 |
| HFF-1 vs. A-253 | 9.989 | -5.155 to 25.13 | No | ns | 0.3001 |
| HFF-1 vs. UCSF-OT | 12.71 | -4.003 to 29.43 | No | ns | 0.1894 |
| HFF-1 vs. FaDu | 4.012 | -12.71 to 20.73 | No | ns | 0.9366 |
| Row 4 | | | | | |
| HFF-1 vs. CAL27 | 32.23 | 17.99 to 46.48 | Yes | **** | <0.0001 |
| HFF-1 vs. A-253 | 26.24 | 11.99 to 40.48 | Yes | **** | <0.0001 |
| HFF-1 vs. UCSF-OT | 16.10 | 0.1685 to 32.02 | Yes | * | 0.0467 |
| HFF-1 vs. FaDu | 13.60 | -2.329 to 29.52 | No | ns | 0.1190 |
| Row 5 | | | | | |
| HFF-1 vs. CAL27 | 36.98 | 22.74 to 51.23 | Yes | **** | <0.0001 |
| HFF-1 vs. A-253 | 35.10 | 20.85 to 49.35 | Yes | **** | <0.0001 |
| HFF-1 vs. UCSF-OT | 25.20 | 9.275 to 41.13 | Yes | *** | 0.0005 |
| HFF-1 vs. FaDu | 18.14 | 2.216 to 34.07 | Yes | * | 0.0195 |
| Row 6 | | | | | |
| HFF-1 vs. CAL27 | 38.25 | 24.00 to 52.50 | Yes | **** | <0.0001 |
| HFF-1 vs. A-253 | 42.29 | 28.04 to 56.53 | Yes | **** | <0.0001 |
| HFF-1 vs. UCSF-OT | 24.76 | 8.834 to 40.69 | Yes | *** | 0.0006 |
| HFF-1 vs. FaDu | 17.05 | 1.120 to 32.97 | Yes | * | 0.0315 |

**c**

| Dunnett's multiple comparisons test | Predicted (LS) mean diff. | 95.00% CI of diff. | Significant? | Summary | Adjusted P Value |
|---|---|---|---|---|---|
| Row 1 | | | | | |
| HFF-1 vs. CAL27 | 1.667e-006 | -19.91 to 19.91 | No | ns | >0.9999 |
| HFF-1 vs. A-253 | -3.333e-006 | -17.81 to 17.81 | No | ns | >0.9999 |
| HFF-1 vs. UCSF-OT | 3.333e-006 | -17.81 to 17.81 | No | ns | >0.9999 |
| HFF-1 vs. FaDu | 6.667e-006 | -19.91 to 19.91 | No | ns | >0.9999 |
| Row 2 | | | | | |
| HFF-1 vs. CAL27 | 31.74 | 11.83 to 51.65 | Yes | *** | 0.0004 |
| HFF-1 vs. A-253 | 28.44 | 10.63 to 46.25 | Yes | *** | 0.0004 |
| HFF-1 vs. UCSF-OT | 23.08 | 5.272 to 40.89 | Yes | ** | 0.0060 |
| HFF-1 vs. FaDu | 5.939 | -13.97 to 25.85 | No | ns | 0.8829 |
| Row 3 | | | | | |
| HFF-1 vs. CAL27 | 28.58 | 8.667 to 48.49 | Yes | ** | 0.0019 |
| HFF-1 vs. A-253 | 27.62 | 9.814 to 45.44 | Yes | *** | 0.0007 |
| HFF-1 vs. UCSF-OT | 24.68 | 6.872 to 42.49 | Yes | ** | 0.0029 |
| HFF-1 vs. FaDu | 25.91 | 5.992 to 45.82 | Yes | ** | 0.0057 |
| Row 4 | | | | | |
| HFF-1 vs. CAL27 | 7.523 | -13.38 to 28.43 | No | ns | 0.7851 |
| HFF-1 vs. A-253 | 8.390 | -10.54 to 27.32 | No | ns | 0.6440 |
| HFF-1 vs. UCSF-OT | 6.704 | -12.23 to 25.64 | No | ns | 0.7943 |
| HFF-1 vs. FaDu | 11.64 | -9.258 to 32.55 | No | ns | 0.4486 |
| Row 5 | | | | | |
| HFF-1 vs. CAL27 | 18.72 | -1.634 to 39.07 | No | ns | 0.0814 |
| HFF-1 vs. A-253 | 7.589 | -10.72 to 25.90 | No | ns | 0.7021 |
| HFF-1 vs. UCSF-OT | 12.38 | -5.931 to 30.70 | No | ns | 0.2834 |
| HFF-1 vs. FaDu | 13.49 | -6.862 to 33.85 | No | ns | 0.3001 |
| Row 6 | | | | | |
| HFF-1 vs. CAL27 | 12.49 | -7.425 to 32.40 | No | ns | 0.3528 |
| HFF-1 vs. A-253 | 5.202 | -12.61 to 23.01 | No | ns | 0.8902 |
| HFF-1 vs. UCSF-OT | 9.766 | -8.045 to 27.58 | No | ns | 0.4756 |
| HFF-1 vs. FaDu | 10.07 | -9.840 to 29.99 | No | ns | 0.5481 |

**d**

| Dunnett's multiple comparisons test | Mean Diff. | 95.00% CI of diff. | Significant? | Summary | Adjusted P Value |
|---|---|---|---|---|---|
| Row 1 | | | | | |
| HFF-1 vs. CAL27 | 1.111e-007 | -22.15 to 22.15 | No | ns | >0.9999 |
| HFF-1 vs. A-253 | 9.000e-006 | -22.15 to 22.15 | No | ns | >0.9999 |
| HFF-1 vs. UCSF-OT | 5.667e-006 | -22.15 to 22.15 | No | ns | >0.9999 |
| HFF-1 vs. FaDu | 1.222e-006 | -22.15 to 22.15 | No | ns | >0.9999 |
| Row 2 | | | | | |
| HFF-1 vs. CAL27 | 8.258 | -13.89 to 30.41 | No | ns | 0.7679 |
| HFF-1 vs. A-253 | 8.087 | -14.07 to 30.24 | No | ns | 0.7803 |
| HFF-1 vs. UCSF-OT | -2.842 | -24.99 to 19.31 | No | ns | 0.9936 |
| HFF-1 vs. FaDu | 3.221 | -18.93 to 25.37 | No | ns | 0.9896 |
| Row 3 | | | | | |
| HFF-1 vs. CAL27 | 1.566 | -20.59 to 23.72 | No | ns | 0.9991 |
| HFF-1 vs. A-253 | 6.778 | -15.37 to 28.93 | No | ns | 0.8664 |
| HFF-1 vs. UCSF-OT | -7.233 | -29.39 to 14.92 | No | ns | 0.8384 |
| HFF-1 vs. FaDu | 14.51 | -7.646 to 36.66 | No | ns | 0.3084 |
| Row 4 | | | | | |
| HFF-1 vs. CAL27 | 20.44 | -1.711 to 42.59 | No | ns | 0.0798 |
| HFF-1 vs. A-253 | 11.08 | -11.07 to 33.23 | No | ns | 0.5476 |
| HFF-1 vs. UCSF-OT | 6.055 | -16.10 to 28.21 | No | ns | 0.9055 |
| HFF-1 vs. FaDu | 14.85 | -7.303 to 37.00 | No | ns | 0.2886 |
| Row 5 | | | | | |
| HFF-1 vs. CAL27 | 14.31 | -7.840 to 36.47 | No | ns | 0.3199 |
| HFF-1 vs. A-253 | 10.51 | -11.65 to 32.66 | No | ns | 0.5929 |
| HFF-1 vs. UCSF-OT | 8.847 | -13.31 to 31.00 | No | ns | 0.7237 |
| HFF-1 vs. FaDu | 1.207 | -20.95 to 23.36 | No | ns | 0.9998 |
| Row 6 | | | | | |
| HFF-1 vs. CAL27 | 19.37 | -2.786 to 41.52 | No | ns | 0.1052 |
| HFF-1 vs. A-253 | 20.98 | -1.176 to 43.13 | No | ns | 0.0692 |
| HFF-1 vs. UCSF-OT | 19.76 | -2.393 to 41.91 | No | ns | 0.0953 |
| HFF-1 vs. FaDu | 0.4882 | -21.66 to 22.64 | No | ns | >0.9999 |

**Fig. S3** Statistical analysis of MTT in A-253, CAL27, UCSF-OT, FaDu and HFF-1 exposed to (**a**) fisetin, (**b**) MKT218, (**c**) MKT257 and (**d**) MKT261 for 24 h. Row 1-6 describes consecutively 3.125 – 50 µM concentrations of substances. Two-way ANOVA, Dunnet’s multiple comparison test. GraphPad Prism 8.

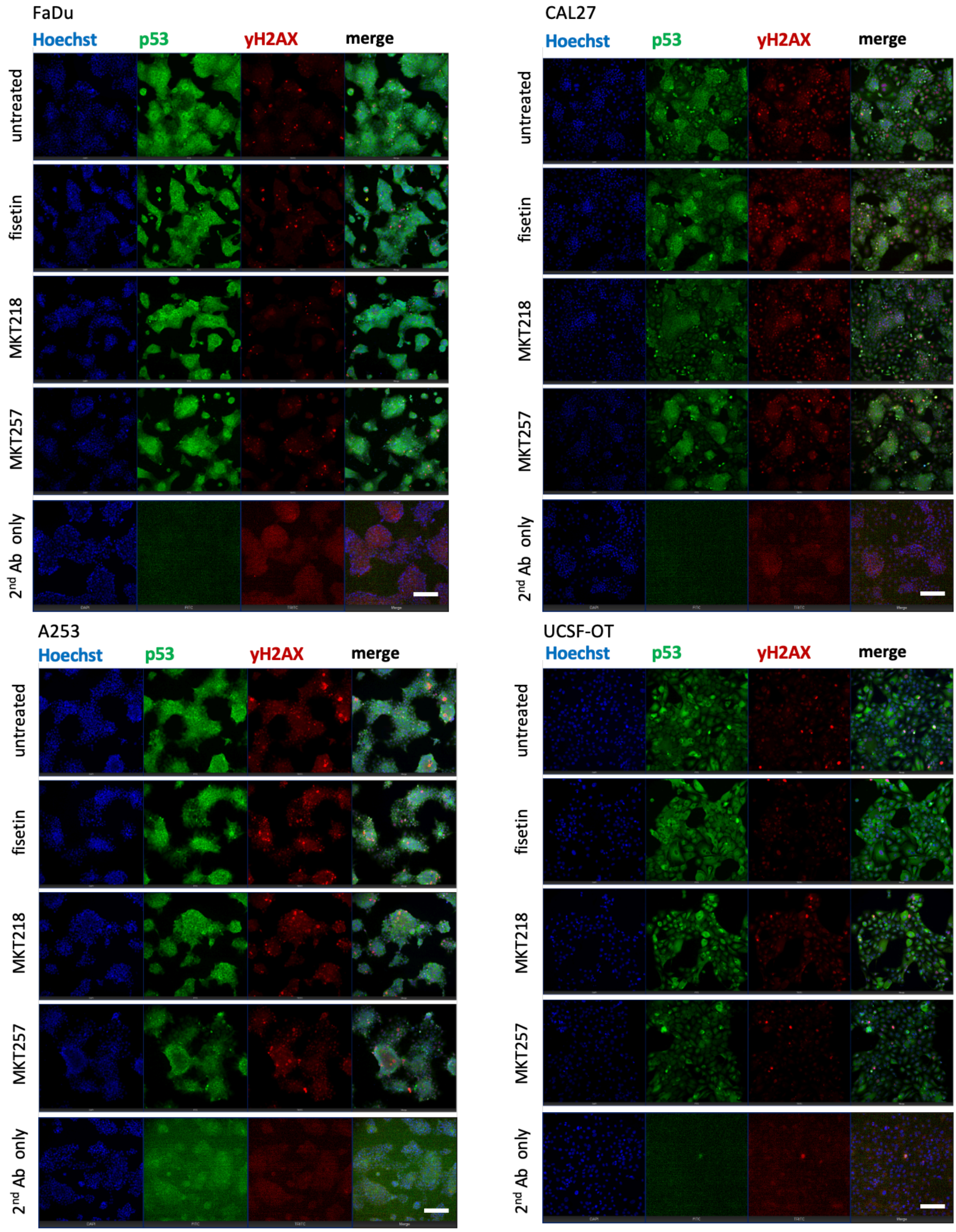

**Fig. S4** Representative images of p53 (green) and γH2AX (red) staining in A-253, CAL27, UCSF-OT and FaDu exposed to fisetin, MKT218, MKT257 (10 μM) for 24 h. Scale bar: 100 μm.

a

| Dunnett's multiple comparisons test | Predicted (LS) mean diff. | 95.00% CI of diff. | Significant? | Summary | Adjusted P Value |
|---|---|---|---|---|---|
| 0h | | | | | |
| Fisetin vs. MKT218 | -1.000 | -3.975 to 1.975 | No | ns | 0.6750 |
| Fisetin vs. MKT257 | 0.1405 | -3.015 to 3.296 | No | ns | 0.9929 |
| 2h | | | | | |
| Fisetin vs. MKT218 | -0.7100 | -3.685 to 2.265 | No | ns | 0.8173 |
| Fisetin vs. MKT257 | -1.555 | -4.710 to 1.601 | No | ns | 0.4426 |
| 4h | | | | | |
| Fisetin vs. MKT218 | 0.1360 | -2.839 to 3.111 | No | ns | 0.9925 |
| Fisetin vs. MKT257 | 0.6130 | -2.543 to 3.769 | No | ns | 0.8741 |
| 6h | | | | | |
| Fisetin vs. MKT218 | 0.5140 | -2.461 to 3.489 | No | ns | 0.8988 |
| Fisetin vs. MKT257 | 2.188 | -0.9675 to 5.344 | No | ns | 0.2148 |
| 8h | | | | | |
| Fisetin vs. MKT218 | -0.5980 | -3.573 to 2.377 | No | ns | 0.8660 |
| Fisetin vs. MKT257 | 3.974 | 0.8180 to 7.129 | Yes | * | 0.0107 |
| 10h | | | | | |
| Fisetin vs. MKT218 | -1.138 | -4.113 to 1.837 | No | ns | 0.6041 |
| Fisetin vs. MKT257 | 4.277 | 1.121 to 7.432 | Yes | ** | 0.0056 |
| 12h | | | | | |
| Fisetin vs. MKT218 | -4.534 | -7.509 to -1.559 | Yes | ** | 0.0017 |
| Fisetin vs. MKT257 | 2.967 | -0.1885 to 6.123 | No | ns | 0.0686 |
| 14h | | | | | |
| Fisetin vs. MKT218 | -6.988 | -9.963 to -4.013 | Yes | **** | <0.0001 |
| Fisetin vs. MKT257 | 2.498 | -0.6580 to 5.653 | No | ns | 0.1409 |
| 16h | | | | | |
| Fisetin vs. MKT218 | -9.286 | -12.26 to -6.311 | Yes | **** | <0.0001 |
| Fisetin vs. MKT257 | 2.755 | -0.4010 to 5.910 | No | ns | 0.0962 |
| 18h | | | | | |
| Fisetin vs. MKT218 | -12.33 | -15.31 to -9.359 | Yes | **** | <0.0001 |
| Fisetin vs. MKT257 | 2.697 | -0.4590 to 5.852 | No | ns | 0.1051 |
| 20h | | | | | |
| Fisetin vs. MKT218 | -12.77 | -15.74 to -9.793 | Yes | **** | <0.0001 |
| Fisetin vs. MKT257 | 2.843 | -0.3130 to 5.998 | No | ns | 0.0838 |
| 22h | | | | | |
| Fisetin vs. MKT218 | -15.62 | -18.60 to -12.64 | Yes | **** | <0.0001 |
| Fisetin vs. MKT257 | 2.720 | -0.4360 to 5.875 | No | ns | 0.1015 |
| 24h | | | | | |
| Fisetin vs. MKT218 | -16.52 | -19.49 to -13.54 | Yes | **** | <0.0001 |
| Fisetin vs. MKT257 | 2.096 | -1.060 to 5.251 | No | ns | 0.2415 |

b

| Dunnett's multiple comparisons test | Mean Diff. | 95.00% CI of diff. | Significant? | Summary | Adjusted P Value |
|---|---|---|---|---|---|
| 0h | | | | | |
| Fisetin vs. MKT218 | -0.3460 | -3.416 to 2.724 | No | ns | 0.9548 |
| Fisetin vs. MKT257 | -0.2600 | -3.330 to 2.810 | No | ns | 0.9741 |
| 2h | | | | | |
| Fisetin vs. MKT218 | -1.236 | -4.306 to 1.834 | No | ns | 0.5710 |
| Fisetin vs. MKT257 | -0.8500 | -3.920 to 2.220 | No | ns | 0.7614 |
| 4h | | | | | |
| Fisetin vs. MKT218 | -0.3920 | -3.462 to 2.678 | No | ns | 0.9424 |
| Fisetin vs. MKT257 | -0.3620 | -3.432 to 2.708 | No | ns | 0.9506 |
| 6h | | | | | |
| Fisetin vs. MKT218 | -0.8960 | -3.966 to 2.174 | No | ns | 0.7393 |
| Fisetin vs. MKT257 | -0.2220 | -3.292 to 2.848 | No | ns | 0.9811 |
| 8h | | | | | |
| Fisetin vs. MKT218 | -1.462 | -4.532 to 1.608 | No | ns | 0.4633 |
| Fisetin vs. MKT257 | -0.1540 | -3.224 to 2.916 | No | ns | 0.9908 |
| 10h | | | | | |
| Fisetin vs. MKT218 | -2.078 | -5.148 to 0.9924 | No | ns | 0.2287 |
| Fisetin vs. MKT257 | -0.3660 | -3.436 to 2.704 | No | ns | 0.9496 |
| 12h | | | | | |
| Fisetin vs. MKT218 | -4.504 | -7.574 to -1.434 | Yes | ** | 0.0026 |
| Fisetin vs. MKT257 | -0.7420 | -3.812 to 2.328 | No | ns | 0.8113 |
| 14h | | | | | |
| Fisetin vs. MKT218 | -6.196 | -9.266 to -3.126 | Yes | **** | <0.0001 |
| Fisetin vs. MKT257 | -0.8360 | -3.906 to 2.234 | No | ns | 0.7680 |
| 16h | | | | | |
| Fisetin vs. MKT218 | -7.512 | -10.58 to -4.442 | Yes | **** | <0.0001 |
| Fisetin vs. MKT257 | -2.012 | -5.082 to 1.058 | No | ns | 0.2488 |
| 18h | | | | | |
| Fisetin vs. MKT218 | -8.118 | -11.19 to -5.048 | Yes | **** | <0.0001 |
| Fisetin vs. MKT257 | -2.228 | -5.298 to 0.8424 | No | ns | 0.1872 |
| 20h | | | | | |
| Fisetin vs. MKT218 | -8.610 | -11.68 to -5.540 | Yes | **** | <0.0001 |
| Fisetin vs. MKT257 | -1.604 | -4.674 to 1.466 | No | ns | 0.4004 |
| 22h | | | | | |
| Fisetin vs. MKT218 | -8.320 | -11.39 to -5.250 | Yes | **** | <0.0001 |
| Fisetin vs. MKT257 | -2.984 | -6.054 to 0.08638 | No | ns | 0.0581 |
| 24h | | | | | |
| Fisetin vs. MKT218 | -9.752 | -12.82 to -6.682 | Yes | **** | <0.0001 |
| Fisetin vs. MKT257 | -4.136 | -7.206 to -1.066 | Yes | ** | 0.0059 |

**Fig. S5** Statistical analysis of timelapses. Percentage of dead cell level in (**a**) A-253 and **(b)** CAL27 exposed to fisetin, MKT218 and MKT257 (10 µM) for 24 h. Two-way ANOVA, Dunnet's multiple comparison test. GraphPad Prism 8.

**a**

| Dunnett's multiple comparisons test | Mean Diff. | 95.00% CI of diff. | Significant? | Summary | Adjusted P Value |
|---|---|---|---|---|---|
| 0h | | | | | |
| Fisetin vs. MKT218 | -1.976 | -8.379 to 4.427 | No | ns | 0.7144 |
| Fisetin vs. MKT257 | -0.8460 | -7.249 to 5.557 | No | ns | 0.9385 |
| 2h | | | | | |
| Fisetin vs. MKT218 | -2.170 | -8.573 to 4.233 | No | ns | 0.6682 |
| Fisetin vs. MKT257 | -1.004 | -7.407 to 5.399 | No | ns | 0.9147 |
| 4h | | | | | |
| Fisetin vs. MKT218 | -3.020 | -9.423 to 3.383 | No | ns | 0.4696 |
| Fisetin vs. MKT257 | -1.838 | -8.241 to 4.565 | No | ns | 0.7464 |
| 6h | | | | | |
| Fisetin vs. MKT218 | -6.164 | -12.57 to 0.2391 | No | ns | 0.0610 |
| Fisetin vs. MKT257 | -2.094 | -8.497 to 4.309 | No | ns | 0.6863 |
| 8h | | | | | |
| Fisetin vs. MKT218 | -7.106 | -13.51 to -0.7029 | Yes | * | 0.0269 |
| Fisetin vs. MKT257 | -2.566 | -8.969 to 3.837 | No | ns | 0.5737 |
| 10h | | | | | |
| Fisetin vs. MKT218 | -12.51 | -18.91 to -6.103 | Yes | **** | <0.0001 |
| Fisetin vs. MKT257 | -2.744 | -9.147 to 3.659 | No | ns | 0.5321 |
| 12h | | | | | |
| Fisetin vs. MKT218 | -15.20 | -21.61 to -8.801 | Yes | **** | <0.0001 |
| Fisetin vs. MKT257 | -0.3540 | -6.757 to 6.049 | No | ns | 0.9889 |
| 14h | | | | | |
| Fisetin vs. MKT218 | -20.52 | -26.92 to -14.11 | Yes | **** | <0.0001 |
| Fisetin vs. MKT257 | -2.170 | -8.573 to 4.233 | No | ns | 0.6682 |
| 16h | | | | | |
| Fisetin vs. MKT218 | -23.12 | -29.53 to -16.72 | Yes | **** | <0.0001 |
| Fisetin vs. MKT257 | -2.442 | -8.845 to 3.961 | No | ns | 0.6031 |
| 18h | | | | | |
| Fisetin vs. MKT218 | -25.09 | -31.50 to -18.69 | Yes | **** | <0.0001 |
| Fisetin vs. MKT257 | -1.216 | -7.619 to 5.187 | No | ns | 0.8779 |
| 20h | | | | | |
| Fisetin vs. MKT218 | -29.98 | -36.38 to -23.57 | Yes | **** | <0.0001 |
| Fisetin vs. MKT257 | -1.994 | -8.397 to 4.409 | No | ns | 0.7101 |
| 22h | | | | | |
| Fisetin vs. MKT218 | -29.77 | -36.17 to -23.36 | Yes | **** | <0.0001 |
| Fisetin vs. MKT257 | -1.844 | -8.247 to 4.559 | No | ns | 0.7450 |
| 24h | | | | | |
| Fisetin vs. MKT218 | -31.35 | -37.75 to -24.94 | Yes | **** | <0.0001 |
| Fisetin vs. MKT257 | -2.094 | -8.497 to 4.309 | No | ns | 0.6863 |

**b**

| Dunnett's multiple comparisons test | Predicted (LS) mean diff. | 95.00% CI of diff. | Significant? | Summary | Adjusted P Value |
|---|---|---|---|---|---|
| 0h | | | | | |
| Fisetin vs. MKT218 | -1.885 | -12.34 to 8.569 | No | ns | 0.8857 |
| Fisetin vs. MKT257 | -3.741 | -14.19 to 6.713 | No | ns | 0.6323 |
| 2h | | | | | |
| Fisetin vs. MKT218 | -4.843 | -15.30 to 5.611 | No | ns | 0.4751 |
| Fisetin vs. MKT257 | -8.589 | -19.04 to 1.865 | No | ns | 0.1215 |
| 4h | | | | | |
| Fisetin vs. MKT218 | -6.191 | -16.64 to 4.263 | No | ns | 0.3108 |
| Fisetin vs. MKT257 | -8.821 | -19.27 to 1.633 | No | ns | 0.1096 |
| 6h | | | | | |
| Fisetin vs. MKT218 | -10.62 | -21.08 to -0.1692 | Yes | * | 0.0458 |
| Fisetin vs. MKT257 | -11.50 | -21.95 to -1.043 | Yes | * | 0.0287 |
| 8h | | | | | |
| Fisetin vs. MKT218 | -16.31 | -26.76 to -5.857 | Yes | ** | 0.0013 |
| Fisetin vs. MKT257 | -12.83 | -23.28 to -2.373 | Yes | * | 0.0133 |
| 10h | | | | | |
| Fisetin vs. MKT218 | -21.95 | -32.41 to -11.50 | Yes | **** | <0.0001 |
| Fisetin vs. MKT257 | -16.95 | -27.40 to -6.496 | Yes | *** | 0.0008 |
| 12h | | | | | |
| Fisetin vs. MKT218 | -23.23 | -33.69 to -12.78 | Yes | **** | <0.0001 |
| Fisetin vs. MKT257 | -19.15 | -29.60 to -8.693 | Yes | *** | 0.0001 |
| 14h | | | | | |
| Fisetin vs. MKT218 | -25.09 | -35.55 to -14.64 | Yes | **** | <0.0001 |
| Fisetin vs. MKT257 | -19.57 | -30.02 to -9.116 | Yes | *** | 0.0001 |
| 16h | | | | | |
| Fisetin vs. MKT218 | -25.44 | -35.90 to -14.99 | Yes | **** | <0.0001 |
| Fisetin vs. MKT257 | -19.95 | -30.40 to -9.494 | Yes | **** | <0.0001 |
| 18h | | | | | |
| Fisetin vs. MKT218 | -22.61 | -33.06 to -12.15 | Yes | **** | <0.0001 |
| Fisetin vs. MKT257 | -16.22 | -26.68 to -5.769 | Yes | ** | 0.0014 |
| 20h | | | | | |
| Fisetin vs. MKT218 | -20.70 | -31.15 to -10.24 | Yes | **** | <0.0001 |
| Fisetin vs. MKT257 | -16.09 | -26.55 to -5.639 | Yes | ** | 0.0016 |
| 22h | | | | | |
| Fisetin vs. MKT218 | -17.69 | -28.15 to -7.241 | Yes | *** | 0.0005 |
| Fisetin vs. MKT257 | -12.19 | -22.64 to -1.737 | Yes | * | 0.0194 |
| 24h | | | | | |
| Fisetin vs. MKT218 | -15.85 | -26.30 to -5.397 | Yes | ** | 0.0019 |
| Fisetin vs. MKT257 | -12.97 | -23.42 to -2.515 | Yes | * | 0.0122 |

**Fig. S6** Statistical analysis of timelapses. Percentage of dead cell level in (**a**) UCSF-OT and **(b**) FaDu exposed to fisetin, MKT218 and MKT257 (10 µM) for 24 h. Two-way ANOVA, Dunnet's multiple comparison test. GraphPad Prism 8.

**a**

| Dunnett's multiple comparisons test | Predicted (LS) mean diff. | 95.00% CI of diff. | Significant? | Summary | Adjusted P Value |
|---|---|---|---|---|---|
| 0h | | | | | |
| Fisetin vs. MKT218 | 0.000 | -0.09864 to 0.09864 | No | ns | >0.9999 |
| Fisetin vs. MKT257 | 0.000 | -0.1046 to 0.1046 | No | ns | >0.9999 |
| 2h | | | | | |
| Fisetin vs. MKT218 | -0.05727 | -0.1559 to 0.04138 | No | ns | 0.3302 |
| Fisetin vs. MKT257 | 0.01935 | -0.08528 to 0.1240 | No | ns | 0.8851 |
| 4h | | | | | |
| Fisetin vs. MKT218 | -0.05622 | -0.1549 to 0.04242 | No | ns | 0.3428 |
| Fisetin vs. MKT257 | 0.02402 | -0.08061 to 0.1286 | No | ns | 0.8295 |
| 6h | | | | | |
| Fisetin vs. MKT218 | -0.1065 | -0.2052 to -0.007871 | Yes | * | 0.0320 |
| Fisetin vs. MKT257 | -0.08495 | -0.1896 to 0.01967 | No | ns | 0.1284 |
| 8h | | | | | |
| Fisetin vs. MKT218 | -0.05974 | -0.1584 to 0.03890 | No | ns | 0.3017 |
| Fisetin vs. MKT257 | -0.08472 | -0.1894 to 0.01990 | No | ns | 0.1297 |
| 10h | | | | | |
| Fisetin vs. MKT218 | -0.03354 | -0.1322 to 0.06510 | No | ns | 0.6690 |
| Fisetin vs. MKT257 | -0.08779 | -0.1924 to 0.01684 | No | ns | 0.1131 |
| 12h | | | | | |
| Fisetin vs. MKT218 | 0.001136 | -0.09751 to 0.09978 | No | ns | 0.9995 |
| Fisetin vs. MKT257 | -0.07620 | -0.1808 to 0.02843 | No | ns | 0.1859 |
| 14h | | | | | |
| Fisetin vs. MKT218 | 0.08287 | -0.01577 to 0.1815 | No | ns | 0.1126 |
| Fisetin vs. MKT257 | -0.07571 | -0.1803 to 0.02892 | No | ns | 0.1895 |
| 16h | | | | | |
| Fisetin vs. MKT218 | 0.1514 | 0.05273 to 0.2500 | Yes | ** | 0.0016 |
| Fisetin vs. MKT257 | -0.06370 | -0.1683 to 0.04093 | No | ns | 0.2981 |
| 18h | | | | | |
| Fisetin vs. MKT218 | 0.1933 | 0.09467 to 0.2920 | Yes | **** | <0.0001 |
| Fisetin vs. MKT257 | -0.08664 | -0.1913 to 0.01799 | No | ns | 0.1191 |
| 20h | | | | | |
| Fisetin vs. MKT218 | 0.2169 | 0.1183 to 0.3156 | Yes | **** | <0.0001 |
| Fisetin vs. MKT257 | -0.07475 | -0.1794 to 0.02988 | No | ns | 0.1970 |
| 22h | | | | | |
| Fisetin vs. MKT218 | 0.2967 | 0.1981 to 0.3953 | Yes | **** | <0.0001 |
| Fisetin vs. MKT257 | -0.04161 | -0.1462 to 0.06302 | No | ns | 0.5809 |
| 24h | | | | | |
| Fisetin vs. MKT218 | 0.3264 | 0.2278 to 0.4251 | Yes | **** | <0.0001 |
| Fisetin vs. MKT257 | 0.005313 | -0.09932 to 0.1099 | No | ns | 0.9907 |

**b**

| Dunnett's multiple comparisons test | Mean Diff. | 95.00% CI of diff. | Significant? | Summary | Adjusted P Value |
|---|---|---|---|---|---|
| 0h | | | | | |
| Fisetin vs. MKT218 | -0.003200 | -0.1463 to 0.1399 | No | ns | 0.9982 |
| Fisetin vs. MKT257 | -0.003200 | -0.1463 to 0.1399 | No | ns | 0.9982 |
| 2h | | | | | |
| Fisetin vs. MKT218 | 0.004400 | -0.1387 to 0.1475 | No | ns | 0.9965 |
| Fisetin vs. MKT257 | -0.03980 | -0.1829 to 0.1033 | No | ns | 0.7595 |
| 4h | | | | | |
| Fisetin vs. MKT218 | -0.06620 | -0.2093 to 0.07690 | No | ns | 0.4824 |
| Fisetin vs. MKT257 | -0.1204 | -0.2635 to 0.02270 | No | ns | 0.1116 |
| 6h | | | | | |
| Fisetin vs. MKT218 | -0.08700 | -0.2301 to 0.05610 | No | ns | 0.2976 |
| Fisetin vs. MKT257 | -0.1578 | -0.3009 to -0.01470 | Yes | * | 0.0280 |
| 8h | | | | | |
| Fisetin vs. MKT218 | -0.07060 | -0.2137 to 0.07250 | No | ns | 0.4393 |
| Fisetin vs. MKT257 | -0.2440 | -0.3871 to -0.1009 | Yes | *** | 0.0004 |
| 10h | | | | | |
| Fisetin vs. MKT218 | -0.08460 | -0.2277 to 0.05850 | No | ns | 0.3163 |
| Fisetin vs. MKT257 | -0.2996 | -0.4427 to -0.1565 | Yes | **** | <0.0001 |
| 12h | | | | | |
| Fisetin vs. MKT218 | 0.02180 | -0.1213 to 0.1649 | No | ns | 0.9192 |
| Fisetin vs. MKT257 | -0.3068 | -0.4499 to -0.1637 | Yes | **** | <0.0001 |
| 14h | | | | | |
| Fisetin vs. MKT218 | 0.05400 | -0.08910 to 0.1971 | No | ns | 0.6094 |
| Fisetin vs. MKT257 | -0.3356 | -0.4787 to -0.1925 | Yes | **** | <0.0001 |
| 16h | | | | | |
| Fisetin vs. MKT218 | 0.08860 | -0.05450 to 0.2317 | No | ns | 0.2856 |
| Fisetin vs. MKT257 | -0.3216 | -0.4647 to -0.1785 | Yes | **** | <0.0001 |
| 18h | | | | | |
| Fisetin vs. MKT218 | 0.1650 | 0.02190 to 0.3081 | Yes | * | 0.0208 |
| Fisetin vs. MKT257 | -0.2816 | -0.4247 to -0.1385 | Yes | **** | <0.0001 |
| 20h | | | | | |
| Fisetin vs. MKT218 | 0.1916 | 0.04850 to 0.3347 | Yes | ** | 0.0063 |
| Fisetin vs. MKT257 | -0.3092 | -0.4523 to -0.1661 | Yes | **** | <0.0001 |
| 22h | | | | | |
| Fisetin vs. MKT218 | 0.2158 | 0.07270 to 0.3589 | Yes | ** | 0.0019 |
| Fisetin vs. MKT257 | -0.2822 | -0.4253 to -0.1391 | Yes | **** | <0.0001 |
| 24h | | | | | |
| Fisetin vs. MKT218 | 0.2322 | 0.08910 to 0.3753 | Yes | *** | 0.0008 |
| Fisetin vs. MKT257 | -0.2768 | -0.4199 to -0.1337 | Yes | **** | <0.0001 |

**Fig. S7** Statistical analysis of timelapses. Proliferation ratio in (**a**) A-253 and **(b**) CAL27 exposed to fisetin, MKT218 and MKT257 (10 µM) for 24 h. Two-way ANOVA, Dunnet's multiple comparison test. GraphPad Prism 8.

a

| Dunnett's multiple comparis | Mean Diff. | 95.00% CI of diff. | Significar | Summary | Adjusted P Value |
|---|---|---|---|---|---|
| 0h | | | | | |
| Fisetin vs. MKT218 | 0.000 | -0.2033 to 0.2033 | No | ns | >0.9999 |
| Fisetin vs. MKT257 | 0.000 | -0.2033 to 0.2033 | No | ns | >0.9999 |
| 2h | | | | | |
| Fisetin vs. MKT218 | 0.02480 | -0.1785 to 0.2281 | No | ns | 0.9473 |
| Fisetin vs. MKT257 | 0.001600 | -0.2017 to 0.2049 | No | ns | 0.9997 |
| 4h | | | | | |
| Fisetin vs. MKT218 | 0.03760 | -0.1657 to 0.2409 | No | ns | 0.8837 |
| Fisetin vs. MKT257 | 0.002000 | -0.2013 to 0.2053 | No | ns | 0.9996 |
| 6h | | | | | |
| Fisetin vs. MKT218 | 0.1166 | -0.08672 to 0.3199 | No | ns | 0.3368 |
| Fisetin vs. MKT257 | 0.05500 | -0.1483 to 0.2583 | No | ns | 0.7706 |
| 8h | | | | | |
| Fisetin vs. MKT218 | 0.07800 | -0.1253 to 0.2813 | No | ns | 0.5996 |
| Fisetin vs. MKT257 | 0.02100 | -0.1823 to 0.2243 | No | ns | 0.9619 |
| 10h | | | | | |
| Fisetin vs. MKT218 | 0.1466 | -0.05672 to 0.3499 | No | ns | 0.1909 |
| Fisetin vs. MKT257 | 0.02520 | -0.1781 to 0.2285 | No | ns | 0.9456 |
| 12h | | | | | |
| Fisetin vs. MKT218 | 0.2134 | 0.01008 to 0.4167 | Yes | * | 0.0381 |
| Fisetin vs. MKT257 | 0.01360 | -0.1897 to 0.2169 | No | ns | 0.9838 |
| 14h | | | | | |
| Fisetin vs. MKT218 | 0.2354 | 0.03208 to 0.4387 | Yes | * | 0.0202 |
| Fisetin vs. MKT257 | -0.05100 | -0.2543 to 0.1523 | No | ns | 0.7986 |
| 16h | | | | | |
| Fisetin vs. MKT218 | 0.2466 | 0.04328 to 0.4499 | Yes | * | 0.0144 |
| Fisetin vs. MKT257 | -0.07400 | -0.2773 to 0.1293 | No | ns | 0.6299 |
| 18h | | | | | |
| Fisetin vs. MKT218 | 0.3240 | 0.1207 to 0.5273 | Yes | *** | 0.0010 |
| Fisetin vs. MKT257 | -0.1426 | -0.3459 to 0.06072 | No | ns | 0.2071 |
| 20h | | | | | |
| Fisetin vs. MKT218 | 0.3972 | 0.1939 to 0.6005 | Yes | **** | <0.0001 |
| Fisetin vs. MKT257 | -0.2062 | -0.4095 to -0.002876 | Yes | * | 0.0463 |
| 22h | | | | | |
| Fisetin vs. MKT218 | 0.4284 | 0.2251 to 0.6317 | Yes | **** | <0.0001 |
| Fisetin vs. MKT257 | -0.2262 | -0.4295 to -0.02288 | Yes | * | 0.0265 |
| 24h | | | | | |
| Fisetin vs. MKT218 | 0.5192 | 0.3159 to 0.7225 | Yes | **** | <0.0001 |
| Fisetin vs. MKT257 | -0.1808 | -0.3841 to 0.02252 | No | ns | 0.0886 |

b

| Dunnett's multiple comparisons test | Mean Diff. | 95.00% CI of diff. | Significant? | Summary | Adjusted P Value |
|---|---|---|---|---|---|
| 0h | | | | | |
| Fisetin vs. MKT218 | 0.000 | -0.1603 to 0.1603 | No | ns | >0.9999 |
| Fisetin vs. MKT257 | 0.000 | -0.1603 to 0.1603 | No | ns | >0.9999 |
| 2h | | | | | |
| Fisetin vs. MKT218 | -0.02600 | -0.1863 to 0.1343 | No | ns | 0.9091 |
| Fisetin vs. MKT257 | 0.02580 | -0.1345 to 0.1861 | No | ns | 0.9104 |
| 4h | | | | | |
| Fisetin vs. MKT218 | 0.02740 | -0.1329 to 0.1877 | No | ns | 0.8997 |
| Fisetin vs. MKT257 | 0.03960 | -0.1207 to 0.1999 | No | ns | 0.8040 |
| 6h | | | | | |
| Fisetin vs. MKT218 | 0.06420 | -0.09614 to 0.2245 | No | ns | 0.5743 |
| Fisetin vs. MKT257 | 0.08980 | -0.07054 to 0.2501 | No | ns | 0.3529 |
| 8h | | | | | |
| Fisetin vs. MKT218 | 0.1334 | -0.02694 to 0.2937 | No | ns | 0.1167 |
| Fisetin vs. MKT257 | 0.1388 | -0.02154 to 0.2991 | No | ns | 0.0994 |
| 10h | | | | | |
| Fisetin vs. MKT218 | 0.1900 | 0.02966 to 0.3503 | Yes | * | 0.0171 |
| Fisetin vs. MKT257 | 0.2282 | 0.06786 to 0.3885 | Yes | ** | 0.0035 |
| 12h | | | | | |
| Fisetin vs. MKT218 | 0.2536 | 0.09326 to 0.4139 | Yes | ** | 0.0011 |
| Fisetin vs. MKT257 | 0.2768 | 0.1165 to 0.4371 | Yes | *** | 0.0003 |
| 14h | | | | | |
| Fisetin vs. MKT218 | 0.2606 | 0.1003 to 0.4209 | Yes | *** | 0.0008 |
| Fisetin vs. MKT257 | 0.2756 | 0.1153 to 0.4359 | Yes | *** | 0.0004 |
| 16h | | | | | |
| Fisetin vs. MKT218 | 0.2698 | 0.1095 to 0.4301 | Yes | *** | 0.0005 |
| Fisetin vs. MKT257 | 0.2984 | 0.1381 to 0.4587 | Yes | *** | 0.0001 |
| 18h | | | | | |
| Fisetin vs. MKT218 | 0.2790 | 0.1187 to 0.4393 | Yes | *** | 0.0003 |
| Fisetin vs. MKT257 | 0.2724 | 0.1121 to 0.4327 | Yes | *** | 0.0004 |
| 20h | | | | | |
| Fisetin vs. MKT218 | 0.2950 | 0.1347 to 0.4553 | Yes | *** | 0.0001 |
| Fisetin vs. MKT257 | 0.3314 | 0.1711 to 0.4917 | Yes | **** | <0.0001 |
| 22h | | | | | |
| Fisetin vs. MKT218 | 0.2512 | 0.09086 to 0.4115 | Yes | ** | 0.0012 |
| Fisetin vs. MKT257 | 0.2854 | 0.1251 to 0.4457 | Yes | *** | 0.0002 |
| 24h | | | | | |
| Fisetin vs. MKT218 | 0.2512 | 0.09086 to 0.4115 | Yes | ** | 0.0012 |
| Fisetin vs. MKT257 | 0.3610 | 0.2007 to 0.5213 | Yes | **** | <0.0001 |

**Fig. S8** Statistical analysis of timelapses. Percentage of dead cell level in (**a**) UCSF-OT and **(b**) FaDu exposed to fisetin, MKT218 and MKT257 (10 µM) for 24 h. Two-way ANOVA, Dunnet's multiple comparison test. GraphPad Prism 8.

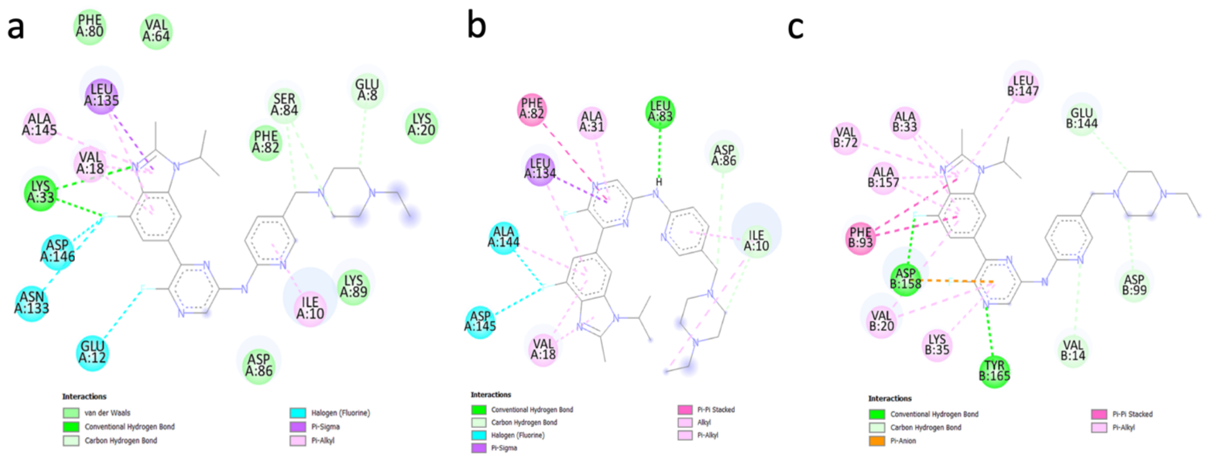

**Fig. S9** The visualization of interaction between Abemaciclib and (**a**) CDK1, (**b**) CDK2, (**c**) CDK4.

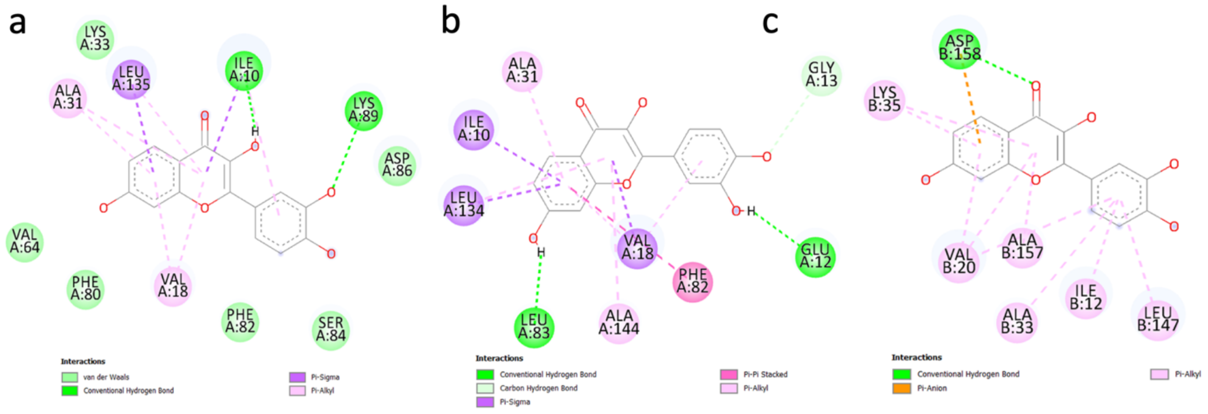

**Fig. S10** The visualization of interaction between fisetin and (**a**) CDK1, (**b**) CDK2, (**c**) CDK4.

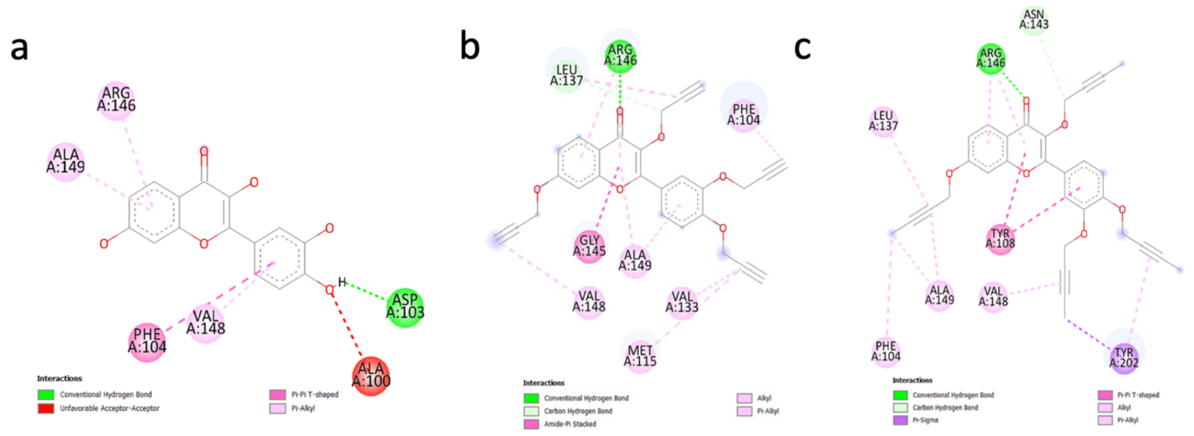

**Fig. S11** Visualization of interaction between Bcl-2 active site and (**a**) fisetin, (**b**) MKT218, (**c**) MKT257.

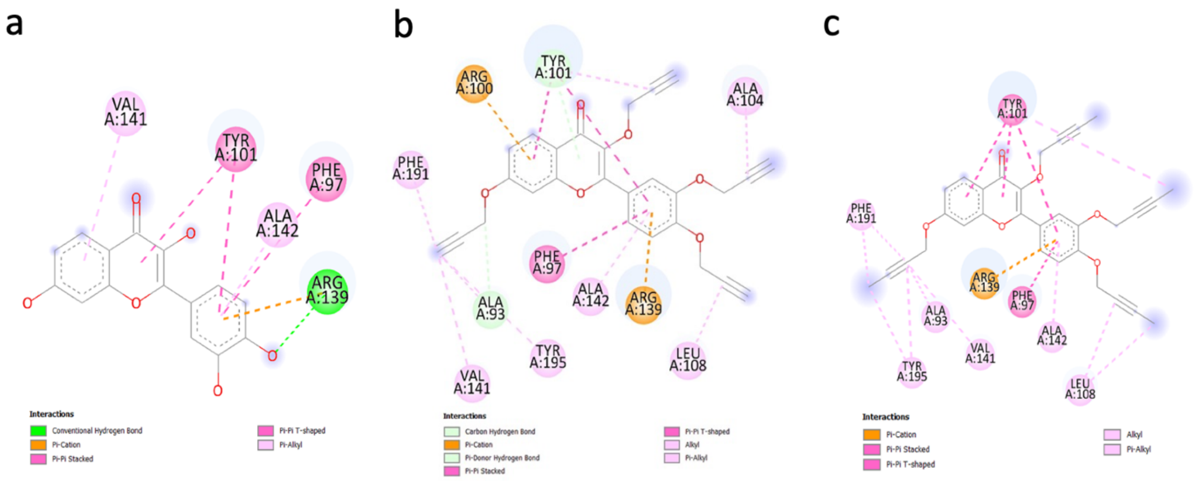

**Fig. S12** Visualization of interaction between Bcl-XL active site and (**a**) fisetin, (**b**) MKT218, (**c**) MKT257.

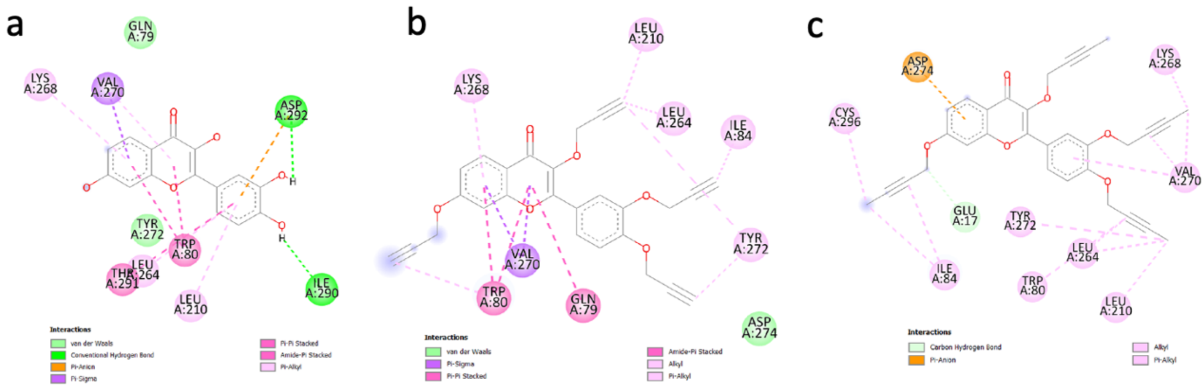

**Fig. S13** Visualization of interaction between AKT1 active site and (**a**) fisetin, (**b**) MKT218, (**c**) MKT257.

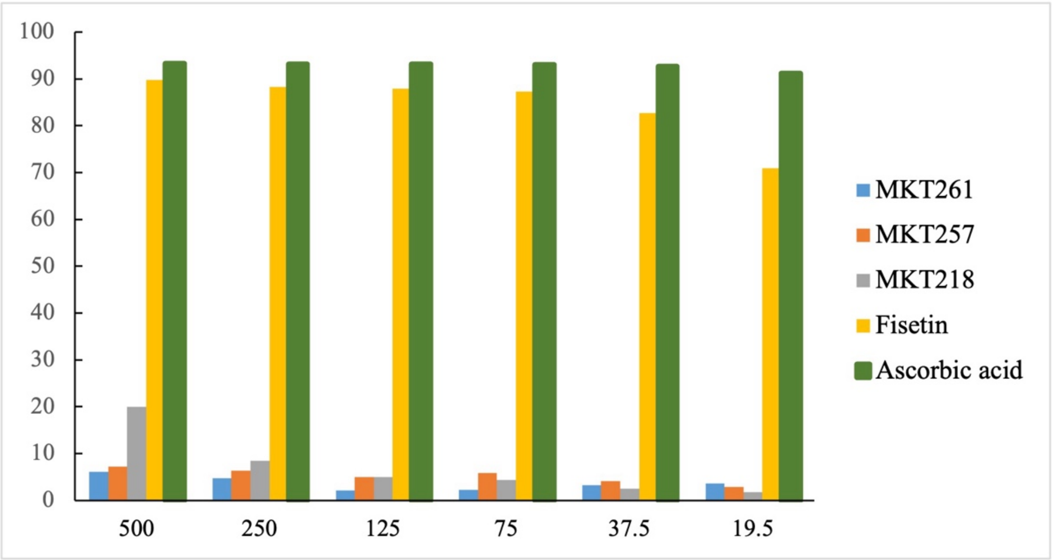

**Fig. S14** Determination of antioxidant DPPH free radical scavenging activity of fisetin, MKT218, MKT257, MKT261 and ascorbic acid in a concentration-dependent manner. N=1.
